# Finding a path: Local search behavior of *Drosophila* larvae

**DOI:** 10.1101/2024.11.21.624685

**Authors:** Jessica Kromp, Tilman Triphan, Andreas S. Thum

**Author notes:** **Correspondence**: Andreas S. Thum, Faculty of Life Sciences, Institute of Biology, Department of Genetics, University of Leipzig, Talstraße 33, 04103 Leipzig, Germany.

## Abstract

Orientation and navigation are essential features of animals living in changing environments. Typically, animals integrate a variety of allothetic and idiothetic cues to achieve their navigational goals. Allothetic cues, such as visual or chemical landmarks from the environment, provide an external frame of reference. In contrast, idiothetic cues are based on internal proprioceptive feedback and internal copies of motor commands.

When *Drosophila* larvae are exposed briefly to a Teflon container holding a food stimulus, they show a characteristic behavior as soon as the container is removed: They briefly crawl away from the detected resource, remain in its vicinity and then return to the area where they experienced the earlier stimulus. We quantified this behavior with respect to the chemosensory nature of the stimulus, starvation time of the larvae, and agarose concentration of the test plate substrate. We conclude that this behavior represents a centered local search. Furthermore, we exclude various external stimuli (vision and taste), which suggests that possibly idiothetic as opposed to allothetic cues have a stronger influence on the larval local search behavior.

In the long term, this behavioral description will enable us to gain insights into the comparability of larval foraging strategies. We also want to investigate whether, despite the simpler organization of the larval brain and the alleged lack of a central complex, a brain region that is important for orientation and navigation in adult Drosophila and other insects, there are common solutions for the brain circuits underlying search behavior.

## Introduction

Navigation and orientation in the environment are indispensable for an animal’s survival. To reach their navigational goals such as food sources and mating partners or to avoid specific places animals rely on allothetic (from the environment) and idiothetic cues (from its own movements) (Bell, 1990; Bures et al., 1998; Mittelstaedt and Mittelstaedt, 2001). Animals often follow visual or olfactory stimuli to find their targets (Zeil, 2023; Zjacic and Scholz, 2022). Bees, living in feature-rich habitats, absolve orientation flights to get familiar with the surrounding environmental features before searching for distant food (Degen et al., 2016; Menzel et al., 1996). In featureless habitats animals can utilize self-motion cues and internal representations to navigate, a process called internal path integration (Muller and Wehner, 1988). Desert ants, for example, live in habitats with few environmental cues; they return home in straight lines after traveling long, winding paths when leaving the nest. This navigation ability is facilitated by the integration of various sensory cues, including celestial and polarization compass cues, along with a pedometer (Muller and Wehner, 1988; Wittlinger et al., 2006).

The fruit fly *Drosophila melanogaster* is a widely used model organism to characterize the neuronal and molecular mechanisms of a multitude of natural behaviors such as locomotion, chemosensory discrimination, courtship, learning and memory, as well as foraging. After consuming a food droplet, adult *Drosophila* perform a specific sequence of repetitive locomotory patterns starting with a pause, a walk away from the stimulus, a reversal, and a return to the previously encountered food source (Corfas et al., 2019; Kim and Dickinson, 2017; Murata et al., 2017; Titova et al., 2023). A similar locomotory pattern was observed in *Drosophila* larvae tested in patchy environments. Depending on the food quality, larvae decreased their velocity, performed higher number of pauses and turned to the center of the patch when reaching its border (Wosniack et al., 2022). Whether larvae perform an adult-like local search after the removal of the food stimulus is not known and was therefore investigated here.

The locomotion of *Drosophila* larvae consists of an alternating sequence of runs and oriented turns. The modulation of locomotion is usually based on allothetic cues such as odors or tastes, which are perceived via the peripheral and pharyngeal nervous system. Olfactory information is received via specific olfactory receptors (ORs) located in 21 olfactory receptor neurons (ORNs) (per body half) within the dorsal organ (Fishilevich et al., 2005; Larsson et al., 2004; Singh and Singh, 1984). In contrast, taste information activates a group of the approximately 55 gustatory receptor neurons (GRNs) (per body half) organized in four external (terminal, ventral, dorsal and labial organ) and four internal head organs (dorsal, ventral, posterior pharyngeal organ, dorsal pharyngeal organ) (Gendre et al., 2004; Python and Stocker, 2002; Richter et al., 2024; Rist and Thum, 2017). When a larva approaches a food source, it modulates its locomotion by making fewer turns, thereby moving in a straighter line. To initiate a turn, the larva scans the local odor gradient by moving its head from side to side, a behavior known as head casting. The larva then performs its next run most often towards the odor source (Gershow et al., 2012; Gomez-Marin et al., 2011).

In the brains of adult flies and many other insects, the central complex (CX) allows the animal to modulate its turn rate and turn direction. In general, the CX consists of four compartments: ellipsoid body (EB), fan-shaped body, protocerebral bridge (PB) and paired noduli (Hanesch et al., 1989). Recently, the asymmetric body is discussed as a fifth compartment (Wolff and Rubin, 2018). Besides its involvement in the orientation toward stable landmarks or the sun, the CX is known to mediate path integration based on internal representations by recalculating the animals’ recent spatial position via self-motion cues (Giraldo et al., 2018; Green et al., 2017; Green et al., 2019; Seelig and Jayaraman, 2015). The current position is established as an activity ‘bump’ within the EB due to the global inhibition of EB-PB-gall-neurons and PB-EB-noduli-neurons and updated by reciprocal exciting interaction of these neurons (Green et al., 2017; Vafidis et al., 2022). However, in larvae, the corresponding neurons are largely undifferentiated, and the few existing parts are non-functional (Farnworth et al., 2020; Riebli et al., 2013). In contrast, it has been shown that the mushroom body (MB), which serves as the integration and memory center, can modulate larval locomotion. Activation of specific MB output neurons prompts the larva to halt and begin head casting while other induces the opposite behavior, causing larvae to suppress turns and move in a straight path (Eichler et al., 2017; Eschbach et al., 2021; Pauls et al., 2010; Saumweber et al., 2018).

To better understand the orientation and foraging behavior of larvae, we have developed a novel behavioral assay. Our findings suggest that upon detecting a chemosensory stimulus, larvae initiate a local search behavior, which continues in darkness for several minutes after the stimulus has been removed. Olfactory and gustatory stimuli, such as apple juice and yeast, trigger local search behavior. Notably, the larval feeding state and the agarose substrate used in the test arena do not significantly influence this behavior. In the long term, the new behavioral paradigm will enable us to identify the neuronal, molecular, and physiological foundations of larval foraging behavior.

## Materials and Methods

### Fly Strains

Flies were reared on standard food medium at 25°C, at a relative humidity of 60-80% and a light/dark cycle of 14/10. Larvae with the *white* mutation (*w^1118^*, Bloomington Stock Centre No.: 3605) were used to determine the parameters for the standard experiment. To test whether the *white* mutation impacts local search behavior, we compared them to wild type Canton-S (*WT-CS*, Bloomington Stock Centre No.: 9515) larvae.

### Recording settings

Larval tracking was performed under red light via the GigE Basler acA1300–60gm NIR camera (version: 106202-22) using the FIM table (Risse et al., 2013) and an adapted recording program that was established in an earlier study of the lab (Schumann and Triphan, 2020). Image sequences were recorded with two frames per second. Infrared light emitting diodes (IR-LEDs) were used for illumination. For an optimal recording of larvae, the recording settings were established to a resolution of 1024 x 1024, an exposure rate of 2000, a gamma of 3.99 and a gain of 1.0. The FIM LED level was set to 6.

### Preparation of Larvae

A spatula tip of food containing six-to-seven-day old larvae were transferred into a Petri dish lid and washed with water. Subsequently, the cleaned animals were collected in a drop of water within a second Petri dish lid. Dependent on the specific starvation time, larvae remain in this drop of water (for one or three hours of starvation) or will be used directly (starvation time of 0 h). Due to the gradual evaporation of water, larvae scheduled to undergo a six-hour period of starvation were transferred to a vial containing water.

### Preparation of chemosensory stimuli

Food odors, serving as chemical stimuli, were freshly prepared each day up to one hour before the start of the initial trial of an experiment and stored in Eppendorf tubes. Five to ten minutes before the experiment started, containers (Teflon, custom-made) were filled with 10 μl of the respective stimuli. Specifically, the following procedures were implemented for each individual stimulus.

#### Apple juice

Containers were freshly prepared before each trial started. To test the effect of various apple juice concentrations (25%, 50%, 75% and 100%), 100% apple juice (standard supermarket quality) was diluted with the respective amount of water.

#### Amyl acetate (AM)

AM (Fluka Cat. No.: 46022; CAS No.: 628-63-7) was diluted with paraffin oil (Fluka Cat. No.: 76235, CAS No.: 8012-95-1) to adjust a concentration of 1:250. The prepared containers were used for up to three consecutive trials.

#### Yeast

Yeast (cube of fresh baker yeast) was weighed and diluted with the respective amount of water to reach the tested yeast concentrations (25%, 50%, 75% and 100%, e.g. for 25%: 0.25 g yeast + 0.75 g water). Notice, due to the high viscosity of the 75% yeast solution, the containers were filled with a pipet tip of this solution. For the 100% yeast a small piece of yeast was placed into the container. Containers were freshly prepared before starting a new trial. Please note that we experienced slight differences in the attractiveness of the yeast container, likely due to variations of the yeast quality and yeast strain obtained from supermarkets. Therefore, we recommend the use of fermented yeast, which did not induce differences in attractiveness.

#### Fermented yeast

Per gram yeast solution (25%, lukewarm water) 0.05 g sucrose was added and stirred. The solution was prepared approximately one hour prior to the initiation of the first trial and stored in a vial sealed with a ceaprene stopper. For the additional yeast-dependent gustatory stimulus, the lid of the container was wetted with the fermented yeast solution. Containers were replaced with new ones before a new trial started. Following a four-hour period, the solution was replaced.

### Preparation of the tracking area

Agarose plates (85mm diameter, Cat. No.: 82.1472, Sarstedt, Nümbrecht), containing the specified agarose concentrations (0.8%, 1.4%, and 2.0%), were freshly prepared daily at least 30 minutes before the start of the initial trial. Agarose layers had a maximum thickness of two millimeters as established by Risse et al. (2013). Approximately 15 min before the experiment up to seven agarose plates were carefully disengaged with a spatula from their Petri dish and transferred to the acrylic plate of the FIM table. In the following, six-centimeter petri dish lids were placed centralized at the agarose plates to define the search area. The inside walls of these lids were roughened with a file to reduce the likelihood of larvae climbing up the lid. In the course of this study these lids were covered with an opaque adhesive film to reduce background noise during tracking. For each experiment a new agarose plate was used.

### Local search paradigm

Experiments started with the transfer of a single larva via a brush to the arena center. With this transfer a five-minute *baseline phase* was initiated during which the larvae could acclimate and explore the search area. Parallel runs were shifted by 30 sec to allow better handling. Subsequently, the container was placed into the arena center. The larvae were given a time window of up to 15 minutes to locate and engage with the container (interaction defined as touching the container with the head, during the *pre-search phase*). Starting with this first container contact, larvae had one minute to investigate the container (*investigation phase*). After that the container was removed, and the larvae were observed for an additional ten minutes during the *search phase* with only the first five minutes being evaluated in the analysis. The following criteria were in addition applied: Firstly, larvae unable to locate the container during the *pre-search phase* or mistakenly perceived as having touched it were excluded from subsequent data analysis. Secondly, interactions where a larva reached the container from the top via the arena lid were not included in the count. In such cases, the larva was given another opportunity to interact with the container via the agarose substrate at the bottom of the search arena. Lastly, larvae that successfully climbed onto the container during the *investigation phase* were gently removed from it and continued to be observed.

### Additional control experiments

#### Empty container experiments

Experiments with an empty container were designed to assess the influence of the container-touch itself on search behavior. The overall protocol remained consistent, with the only alteration being the placement of an empty container, devoid of any stimulus, into the arena.

#### Without container experiments

Experiments without container were performed to indicate whether larvae change their behavior from avoiding the center to preferring the center over time. Therefore, larvae were tracked for the maximum duration of the standard protocol (31 min) without any interruption (neither container placing / removing nor opening of the arena lid). In the subsequent data evaluation, the tracking sequence was artificially structured into the four experimental phases.

#### Naïve larvae experiments

To eliminate the possibility of lingering odor cues causing or influencing centralized search behavior, experiments involving naïve larvae were performed. The experiment started without larvae at the beginning of the *baseline phase*. Subsequently, a container was introduced into the arena for a duration of 7 minutes and 16 seconds (the mean time a larva needed to find the container when using fermented yeast as a chemical stimulus). Afterwards, the container was removed, and a naïve larva was placed in the center of the arena to assess the potential preference for any remaining odor.

### Image processing and tracking of larvae via ImageJ and FIMtrack

Following the recording, the generated image sequences were pre-processed by a self-written ImageJ script (National Institutes of Health, Bethesda, Maryland, http://imagej.nih.gov/ij). Regions of interest (ROIs) covering the single arenas were set manually and its center coordinates measured. The script masked the area surrounding ROIs, calculated a median image of the image sequence and determined the difference between the masked and median images to reduce background noise. Finally, the processed data were saved as an image sequence in tiff format. These processed image sequences were then opened with the FIMtrack software, and the larval positions tracked (tracking settings: ‘Gray Threshold’: 100, ‘Min larval size’: 10, ‘Max larval size’: 100). If the software failed to track the larva or split the larval track several times, the ‘Gray Threshold’ was reduced stepwise by a value of 20. Subsequently, the single tracks of an individual larva were stitched together.

### Data evaluation

Tracking data were analyzed via a self-written MATLAB script (The MathWorks Inc. 2021). In a first step FIMtrack data (Risse et al., 2013), center coordinates and information of the experiment (e.g. concentration of the chemical stimulus, time point of container placing) were loaded. Due to tracking errors, such as reflections or larvae disappearing at the arena edge or container, the detection of larvae failed for some frames (mean=21.7%, varying between 7.7% and 45.8% dependent on the experiment and experimental phase). Missing detections which started and ended near the arena edge (>2.13 cm distance to center) were interpolated via a circular path. Gaps occurring at the end of the experiment were interpolated by repeating the last tracked coordinate. All other gaps were interpolated via a straight line. Stepwise other parameters were calculated based on the tracking coordinates.

In a first step the distance between the larval coordinates and the center was calculated (Pythagoras’s theorem) and converted to real size (Conversion coefficient: 1 px=0.2367 mm). Equally the distance between consecutive coordinates was computed and cumulated to determine the larval track length. Further, the number of revisits was determined. A revisit was counted if the larva’s center of mass crossed the reward zone boarder. The reward zone is defined as a center surrounding area with a radius of 0.80 cm. The radius was calculated by summing the radius of the container (0.35 cm), the median larval size (∼0.35 cm) and 0.10 cm for inconsistencies (e.g. larval size variances, imprecise container placement). Furthermore, the time spent in the reward zone was calculated and the velocity data of the FIMtrack software evaluated.

To identify where differences occurred, we refined the analysis. The search arena was divided into distance categories (0-4 mm, 4-8 mm, 8-12 mm, 12-16 mm, 16-20 mm, 20-24 mm and >24 mm) and the time spent as well as track length moved in each category determined. Further, the mean distances to the arena center within one-minute time intervals were analyzed. Finally, we calculated a search score reflecting the larval behavior with a single parameter. It was calculated by subtracting the time spent in the edge zone (area adjoining the edge up to a distance of 0.35 cm) from the time spent in a search zone between 8 and 16 mm from the center of the arena, divided by the total time analyzed (here: 5 min). The area between the two zones is defined as neutral. Positive values display a search zone preference, negative an edge preference. For the statistical analyzes and visualizations, the data were split into the different experimental phases. Only the first five minutes of the *search phase* were compared to the *baseline*. The padcat function of Jos (https://de.mathworks.com/matlabcentral/fileexchange/22909-padcat) was applied for the data evaluation. Significances were visualized using the adapted sigstar function of Campbell (https://github.com/raacampbell/sigstar).

### Statistics

The statistical analysis was conducted with the MATLAB script. We kept the statistics conservative by only performing non-parametric tests. Differences between *baseline* and *search phase* were evaluated with the two-sample Wilcoxon signed-rank test. To investigate whether the preference score differed from a random behavior the one-sample Wilcoxon signed-rank test was performed. For the comparison of the behavioral control with naïve larvae the Mann-Whitney U test was established. Results are visualized in boxplots, indicating the median as middle line, 25% / 75% quantiles as box boundaries and minimum / maximum performance indices as whiskers. Significance levels were set to *p≤0.05, **p≤0.01 and ***p<0.001. A detailed statistical evaluation can be found in Table S1-S3.

### AI tools usage in writing the manuscript

AI tools (such as DeepL and ChatGPT) were employed for grammar, punctuation, language and translation checks. However, the definition of the research question, the methodological approach, the interpretation of the data, the conclusions drawn from the results, and their presentation in the appropriate scientific context were all carried out solely by the authors, without AI assistance. The authors bear full responsibility for the ethical considerations of the research.

## Results

### A paradigm to analyze local search behavior in *Drosophila* larvae

Investigations in insects, such as desert ants (*Cataglyphis fortis*) or adult *Drosophila*, have shown that they can return to a navigational goal even in the absence of external stimuli (Kim and Dickinson, 2017; Muller and Wehner, 1988; Murata et al., 2017). To determine whether *Drosophila* larvae are capable of similar behavior, we developed the larval local search paradigm (Fig.1A). Following a one-hour period of starvation, individual larvae were positioned in the center of a circular agarose substrate and their movements were recorded in darkness using the FTIR-based imaging method (FIM, Risse et al., 2013). After observing the larvae’s behavior during the initial *baseline phase*, a Teflon container filled with pure apple juice (100%) was introduced at the arena’s center. The larvae had up to 15 minutes to locate and touch the container during the *pre-search phase* (Movie 1). They then had 1 minute to explore the container, followed by 5 minutes of observation after the container was removed (*search phase*). Larvae unable to reach the container within the *pre-search phase* were excluded from the analysis. The visualization of the larval tracks reveals that larvae briefly orientate after their placement before departing from the center of the arena (Fig.1B). Throughout the remaining *baseline phase*, larvae predominantly stayed in proximity to the edges. In the *pre-search phase*, larvae explored the arena by circling around the container or moving directly toward it. Of the 55 tested larvae, 20 interacted with the container and were subjected to further observation. While most larvae remained close to the container, a few had already left during the *investigation phase*. Following the removal of the container, larvae conducted local searches around the previous food spot, with some leaving the center and returning to the edges (Fig.1B).

**Figure 1:**
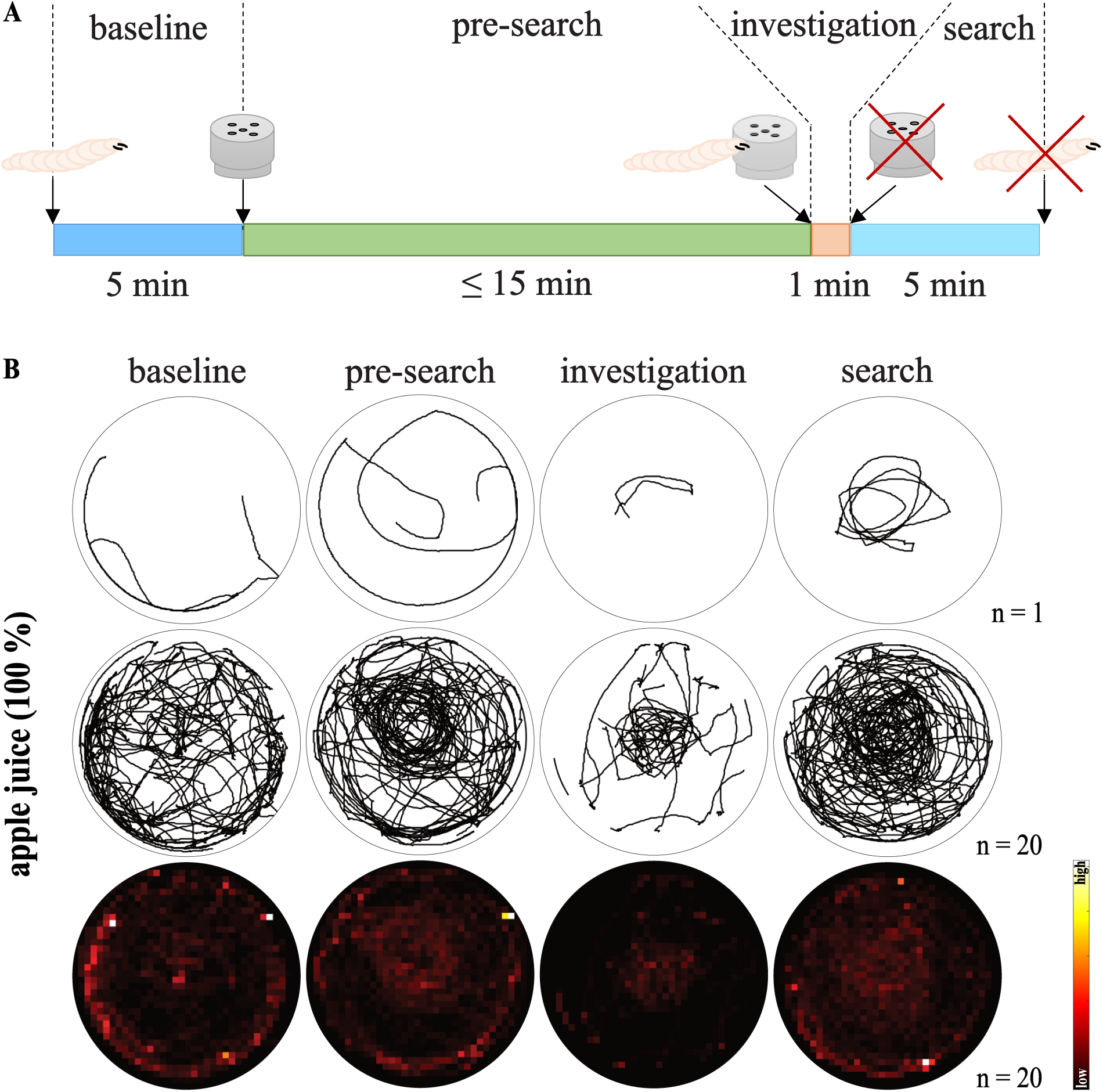
*Drosophila melanogaster* larvae perform a local search. (A) The local search paradigm. Individuals were placed into the center of a search arena (5.8 cm diameter) and had five minutes time to explore it (*baseline phase*). Next, an odor-filled container was placed into the arena center and initiated the *pre-search phase*. If the larva interacted with the container (touched via head) within 15 min a one-minute *investigation phase* started. The *investigation phase* was terminated with the removing of the container. Subsequently, the larvae were observed for five minutes (*search phase*). Larvae that failed to interact with the container were excluded. (B) Walking trajectories/residency plots - apple juice (100%). Columns show the crawling path and residency during the four test phases (*baseline, pre-search, investigation,* and *search*) row-wise for a single larva or the entire test group. Larvae were tested after one-hour starvation time on 0.8% agarose plates.

### A parametric analysis of larval local search behavior

Based on the observed changes in larval behavior, we established a processing pipeline to identify and quantify parameters indicative of a local search. First, we compared the distance of the larvae from the center between the *baseline* and *search phases* (Fig.2A,B). While larvae leave the arena’s center within the first minute of the *baseline phase* and remain, thereafter, close to the edge, they exhibit a more consistent and closer proximity to the center during the *search phase* (Fig.2C). A direct comparison of one-minute time intervals shows that initially larvae start both *phases* at equivalent distances (Fig.2D). However, in subsequent one-minute intervals, with the exception of the fourth, larvae stay closer to the center during the *search phase*. The larvae do not spend more time in a “reward zone” (Fig.2E). However, a more detailed evaluation, by dividing the arena radius into seven distinct distance categories revealed that larvae spent more time within a distance of 8-16 mm from the center and less time within the outermost distance category during the *search phase* (Fig.2F,G). This means that larvae circulated around the previous container position.

**Figure 2:**
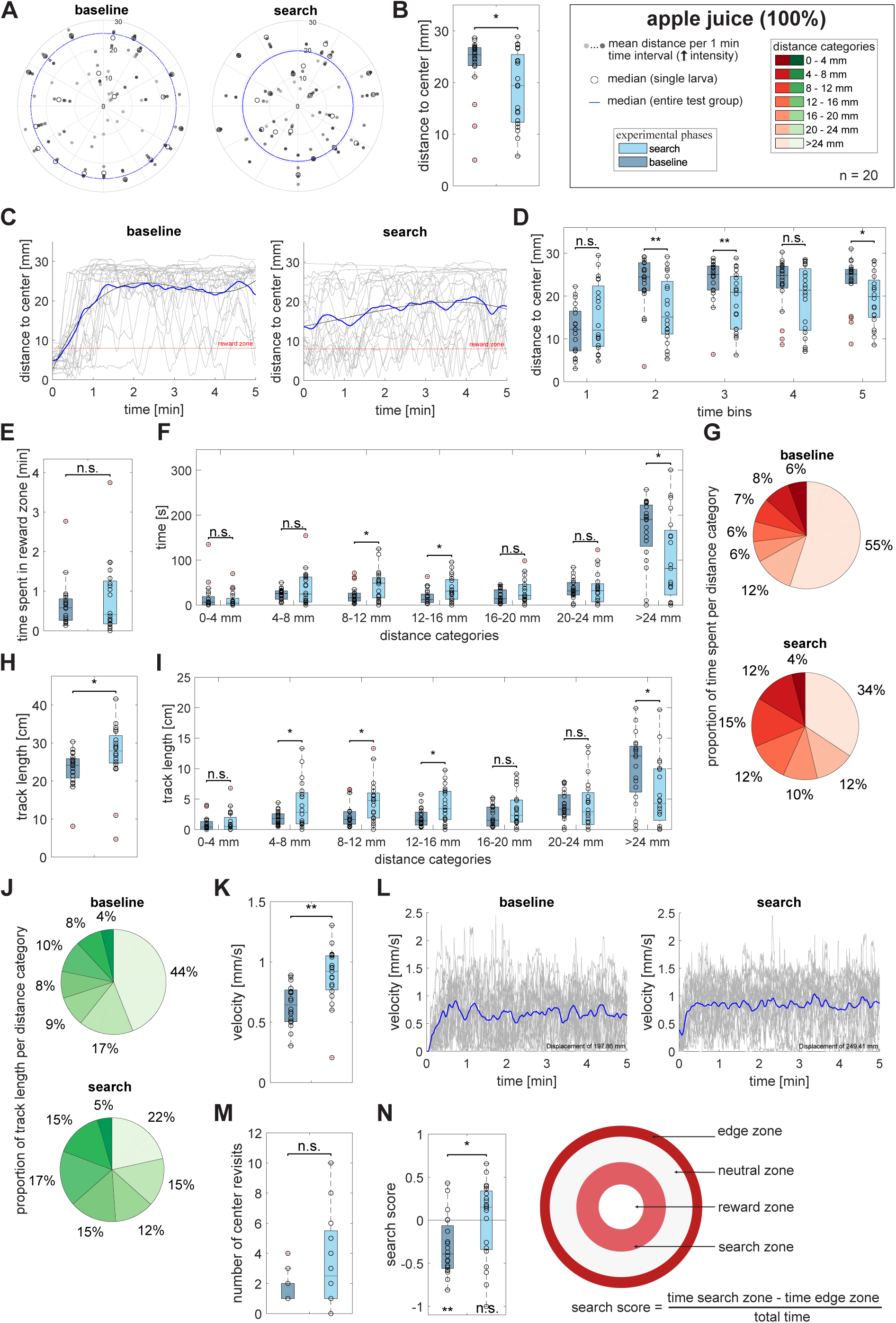
Evaluation of tracking data of larvae exposed to apple juice (100%). (A) Distance to center – polar scatter. The figure displays the individuals mean distances to center within one-minute time intervals, the individual median distance during the entire phase, and the median distance of the entire test group. Individuals are represented by circles on a line extending from the center. (B) Distance to center – boxplot. Compared are the median distances of larvae during the *baseline* and *search phase*. Larvae reduced their distances to center during the *search phase* significantly compared to the *baseline* (p=0.019). (C) Distance to center – progression over time. The graphs display the test group’s mean distance to center (blue line) as well as the individual distances (grey lines) over time, averaged over ten frames. At the beginning of the *baseline* the larvae rapidly left the arena center and remained near the edge while they stayed more centralized during the *search phase*. (D) Distance to center – time intervals. Represented are the distances to center during one-minute time intervals. While larvae did not differ in their distance to center within the *baseline* and *search phase* during the first (p=0.135) and fourth (p=0.126) time interval, they remained closer to the center during the *search phase* for the duration of the second (p=0.004), third (p=0.003) and fifth (p=0.037) time intervals. (E) Time spent in reward zone. The comparison of *baseline* and *search phase* revealed no differences in the time spent within the reward zone (p=0.391). (F) Time spent per distance category compared for *baseline* and *search phase*. Larvae spent more time in a distance of 8-16 mm (8-12 mm: p=0.038, 12-16 mm: p=0.018) and lesser time in the outermost distance category (>24 mm: p=0.014) after the exposure to apple juice. The time spent within the other distance categories was unaffected (0-4 mm: p=0.446, 4-8 mm: p=0.173, 16-20 mm: p=0.117, 20-24 mm: p=0.809). (G) Proportion of time spent per distance category. Pie charts represent the proportion of time spent per distance category counterclockwise from the center (dark red) to the edge (beige). (H) Track length. The plots display the total track length covered during the respective phases. Larvae crawled longer paths after the interaction with the container (p=0.017). (I) Track length per distance category. Shown is the track length covered per distance category compared to the respective phases. Larvae crawled longer paths in a distance of 4-16 mm (4-8 mm: p=0.040, 8-12 mm: p=0.023, 12-16 mm: p=0.015) while lowering the crawled distance in the outermost distance category (>24 mm: p=0.010) after an exposure to apple juice. The track length within the remaining distance categories was unaffected (0-4 mm: p=0.396, 16-20 mm: p=0.145, 20-24 mm: p=0.841). (J) Proportion of track length crawled per distance category. Pie charts represent the proportion of track length crawled per distance category counterclockwise from the center (darkest green) to the edge (lightest green). (K) Velocity – boxplot. The boxplot compares the larval velocity during the respective phases. After the container interaction larvae increased their speed (p=0.001). (L) Velocity - progression over time. The graphs display the test group’s mean velocity (blue line) as well as the individual velocity (grey lines) over time, both averaged over ten frames. (M) Number of center revisits. The boxplot visualizes the number of center revisits during the respective phases. A revisit was counted when a larva crosses the reward zone boundary (inward direction). The number of revisits was unaffected by the container interaction (p=0.092). (N) Search score. The search score displays whether larvae prefer the search (positive values) or the edge (negative values) zone. The results show that larvae preferred the edge during the *baseline* (p=0.003) while behaving neutral during the *search phase* (p=0.852). Larvae reduced their center avoidance significantly after odor presentation (p=0.014). The larvae were tested after one-hour starvation time on 0.8% agarose plates. For the statistical evaluation the one-sample and two-sample Wilcoxon signed-rank test were performed. *p≤0.05, **p≤0.01, ***p<0.001

Despite larvae spending increased time within distinct distance categories, this parameter alone does not define a local search behavior. To broaden our analysis, we compared the track length that larvae crawl during *baseline* and *search phases*. We found that larvae crawled longer distances during the *search phase* than during the *baseline phase* (Fig.2H). Particularly within the 4-16 mm distance range, larvae traversed considerably longer paths following the removal of the container. Conversely, a shorter track length was observed in the outermost distance category (Fig.2I,J). The distances covered within the remaining categories remained unaltered. The increased track length observed during the *search phase* is caused by a higher velocity exhibited by the larvae after the presentation of apple juice (Fig.2K,L). However, the larvae do not revisit the center more often during their *search phase* than during the *baseline phase* (Fig.2M). In characterizing the behavior, we also computed a search score (Fig.2N), which represents the ratio of the time larvae spent in the search zone compared to the time spent in the outermost region, divided by the total duration of the respective phase. As expected, the larvae exposed to apple juice exhibit a search score that is more positive than that of naïve larvae during the initial *baseline phase* (Fig.2N, see Fig.S1 to compare different apple juice concentrations). The combination of all the parameters reveals that, following apple juice presentation, the larvae exhibit extended and more focused movements around the prior container position compared to naïve larvae. Hence, we would like to refer to this locomotion pattern as local search behavior of the larvae in the following sections.

### The nature of the chemical stimulus presented exerts varying effects on the local search behavior

Given that apple juice stimulation can induce local search behavior, we subsequently investigated whether other chemical stimuli could elicit a similar response. In addition to an empty container, we used the olfactory stimuli amyl acetate (AM) and yeast and contrasted them to the results with pure apple juice obtained before (Fig.2 and Fig.3A-F, same data). We focused the analysis on the parameters ‘distance to center’, ‘proportion of time spent per distance category’, ‘proportion of track length crawled per distance category’ and ‘search score’ as these allow for a comprehensive description of the behavior.

**Figure 3:**
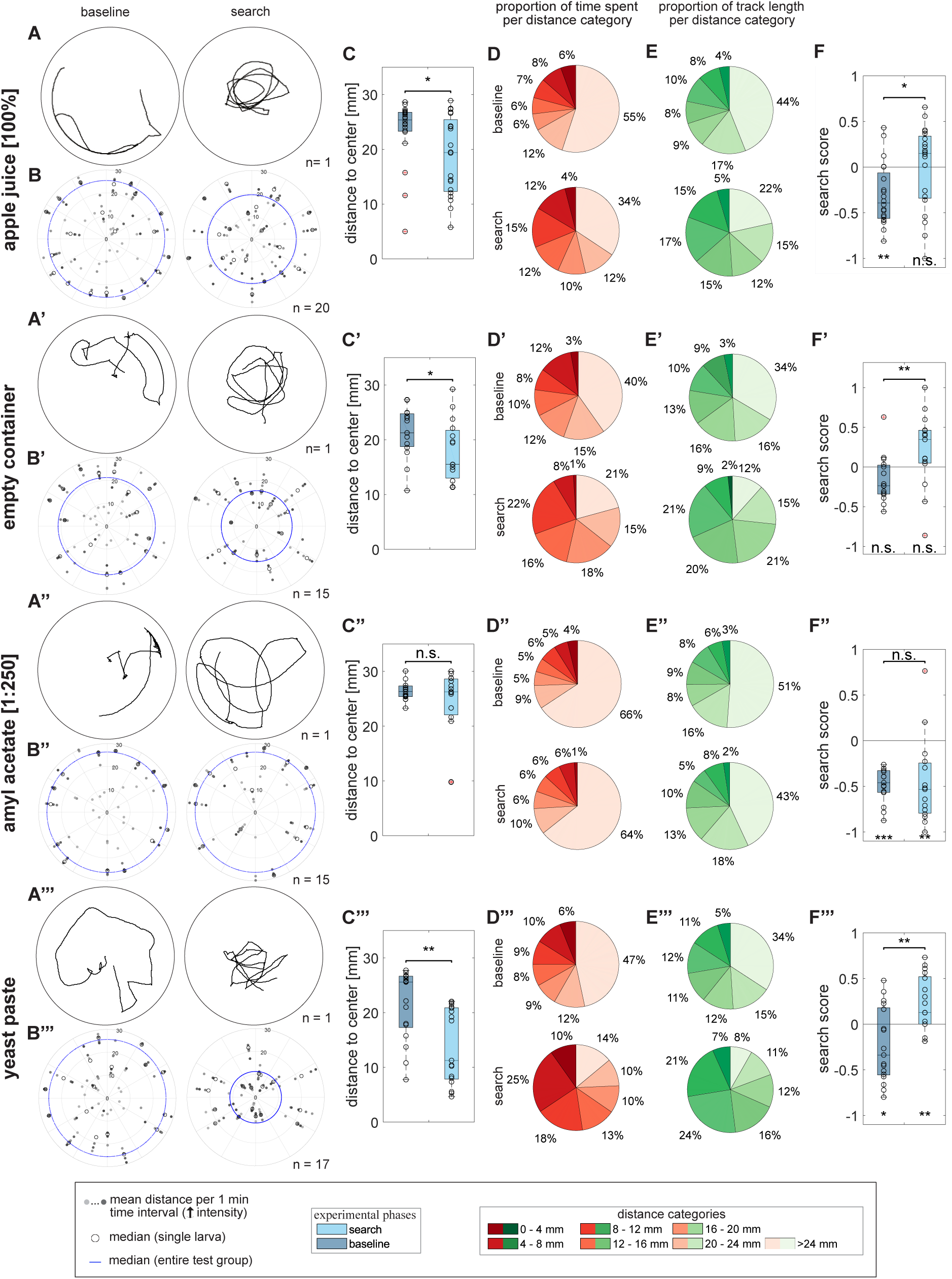
Impact of different chemical stimuli on the local search. Larvae were tested via the local search paradigm to investigate the effect of different chemical stimuli (apple juice, amyl acetate and yeast). Empty container experiments were included to evaluate the influence of the odor container per se. (A) Example track. The plot shows the crawled path before and after the presentation of a stimulus. (B) Distance to center – polar scatter. The figure displays mean distances to center for individuals within one-minute time intervals, the individual median distance during the entire phase, and the median distance of the entire test group. Individual larvae are represented by circles on a line extending from the center. (C) Distance to center – boxplot. Compared are the median distances of larvae during the *baseline* and *search phase*. Larvae exposed to apple juice (C: p=0.019) and yeast (C’’’: p=0.002) as well as an empty container (C’: p=0.022) led to a distance reduction to center while the presentation of AM has no effect (C’’: p=0.934). (D) Proportion of time spent per distance category. Pie charts represent the proportion of time spent in each distance category counterclockwise from the center (dark red) to the edge (beige). (E) Proportion of track length crawled per distance category. Pie charts represent the proportion of track length crawled per distance category counterclockwise from the center (darkest green) to the edge (lightest green). (F) Search score. The results show that each test group, except the empty container group, avoid the center during the *baseline phase* (F: p=0.003; F’: p=0.064; F’’: p<0.001; F’’’: p=0.001). After the container interaction all test groups, except the one exposed to AM, behaved neutrally or preferred the search zone (F: p=0.852; F’: p=0.073; F’’: p=0.005; F’’’: p=0.008). This preference change was significant for larvae exposed to apple juice or yeast (F: p=0.014, F’: p=0.005, F’’: p=0.804, F’’’: p=0.002). The larvae were tested after a starvation time of one-hour on 0.8% agarose plates. To compare other stimuli with apple juice (100%), we present parts of the data of Fig.1 and 2 here again. For the statistical evaluation the one-sample and two-sample Wilcoxon signed-rank test were performed. *p≤0.05, **p≤0.01, ***p<0.001

The results showed that already the empty container interaction triggered local search behavior. After the presentation and removal of the container the larvae remained closer to the center (Fig.3A’-C’), spent less time and moved a shorter route length in distance categories closer to the edge. At the same time, the values of these parameters increased at distances closer to the center, especially in the range from 8-20 mm (Fig.3D’/E’). The interaction with an empty container also increased the search score (Fig.3F’). The introduction of a container filled with AM, a stimulus commonly employed in larval olfactory conditioning (Chen and Gerber, 2014; Scherer et al., 2003; Weber et al., 2023; Widmann et al., 2016), did not evoke local search behavior (Fig.3A’’-F’’, Movie 2), which is surprising given the results for the empty container shown before. Neither the distance to center, nor the search scores were altered. The AM exposed larvae remained in distances closer to the edge of the arena and avoided the search zone significantly. In contrast, yeast paste had a pronounced effect. Larvae reinforced their local search behavior (Fig.3A’’’,Fig.3F’’’) reflected by a stronger reduction of their distance to center (Fig.3B’’’/C’’’) as well as an extended time spent and track length crawled within distances close to the center (Fig.3D’’’/E’’’). Based on the more stable and enhanced effects to induce larval search behavior, we opted to conduct the subsequent experiments using yeast paste.

### Yeast induces local search behavior, regardless of its concentration

Next, we tested how four different yeast concentrations - 25%, 50%, 75% and 100% - trigger local search behavior (Fig.4). The results display a similar outcome for all concentrations. At the onset of the experiment, the larvae departed from the arena’s center and positioned themselves closer to the edge. In contrast, during the *search phase*, they consistently traversed distances near the center, circling around the location where the yeast-filled container had been presented previously (Fig.4A-A’’’,B-B’’’,C-C’’’, Movie 3; left). A more detailed analysis unveiled for all four groups a rise in duration and distance covered within the center-proximate distance categories while markedly decreasing in the outermost distance category (Fig.4D-D’’’,E-E’’’). All four groups showed a significant increase in their search scores (Fig.4F-F’’’). Given these similarities we decided to continue the experiments with a yeast concentration of 25%. This is the weakest tested chemical stimulation, and the larvae locate the container swiftly and frequently in the *pre-search phase* (Table 1).

**Figure 4:**
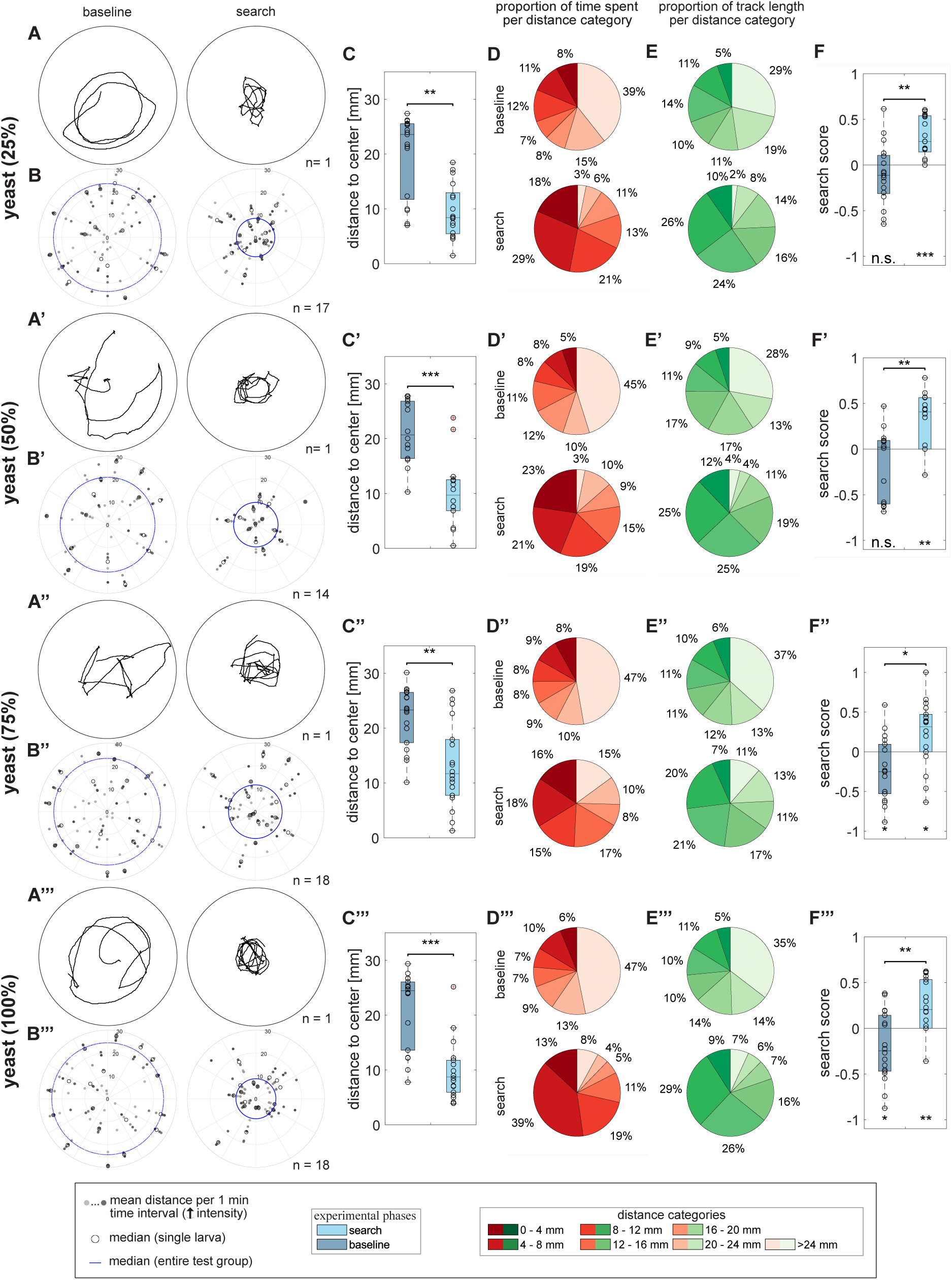
Influence of the yeast on the search behavior. Larvae were tested via the local search paradigm to examine the effect of different yeast concentrations 25%, 50%, 75% and 100%. (A) Example track. The plot shows the crawled path before and after the presentation of yeast. (B) Distance to center – polar scatter. Displayed are individual mean distances to center within one-minute time intervals, individual median distance during the entire phase and median distance of the entire test group. Individuals are represented by circles on a line extending from the center. (C) Distance to center – boxplot. Compared are the median distances of larvae during the respective phases. All test groups reduced their distance to center significantly after the yeast presentation (C: p=0.002, C’: p<0.001, C’’: p=0.006, C’’’: p<0.001). (D) Proportion of time spent per distance category. Pie charts represent the proportion of time spent per distance category counterclockwise from the center (dark red) to the edge (beige). (E) Proportion of track length crawled per distance category. Pie charts represent the proportionw of track length crawled per distance category counterclockwise from the center (darkest green) to the edge (lightest green). (F) Search score. All larvae avoided or behaved neutrally during the *baseline phase* and preferred the search zone after the yeast presentation (F: p_base_=0.210, p_search_<0.001; F’: p_base_=0.021, p_search_=0.003; F’’: p_base_=0.025, p_search_=0.049; F’’’: p_base_=0.039, p_search_=0.003). All test groups increased their search score significantly (F: p=0.003, F’: p=0.005, F’’: p=0.018, F’’’: p=0.001). All larvae were tested after one-hour starvation time on 0.8% agarose plates. For the statistical evaluation the one-sample and two-sample Wilcoxon signed-rank test were performed. *p≤0.05, **p≤0.01, ***p<0.001

**Table 1:**
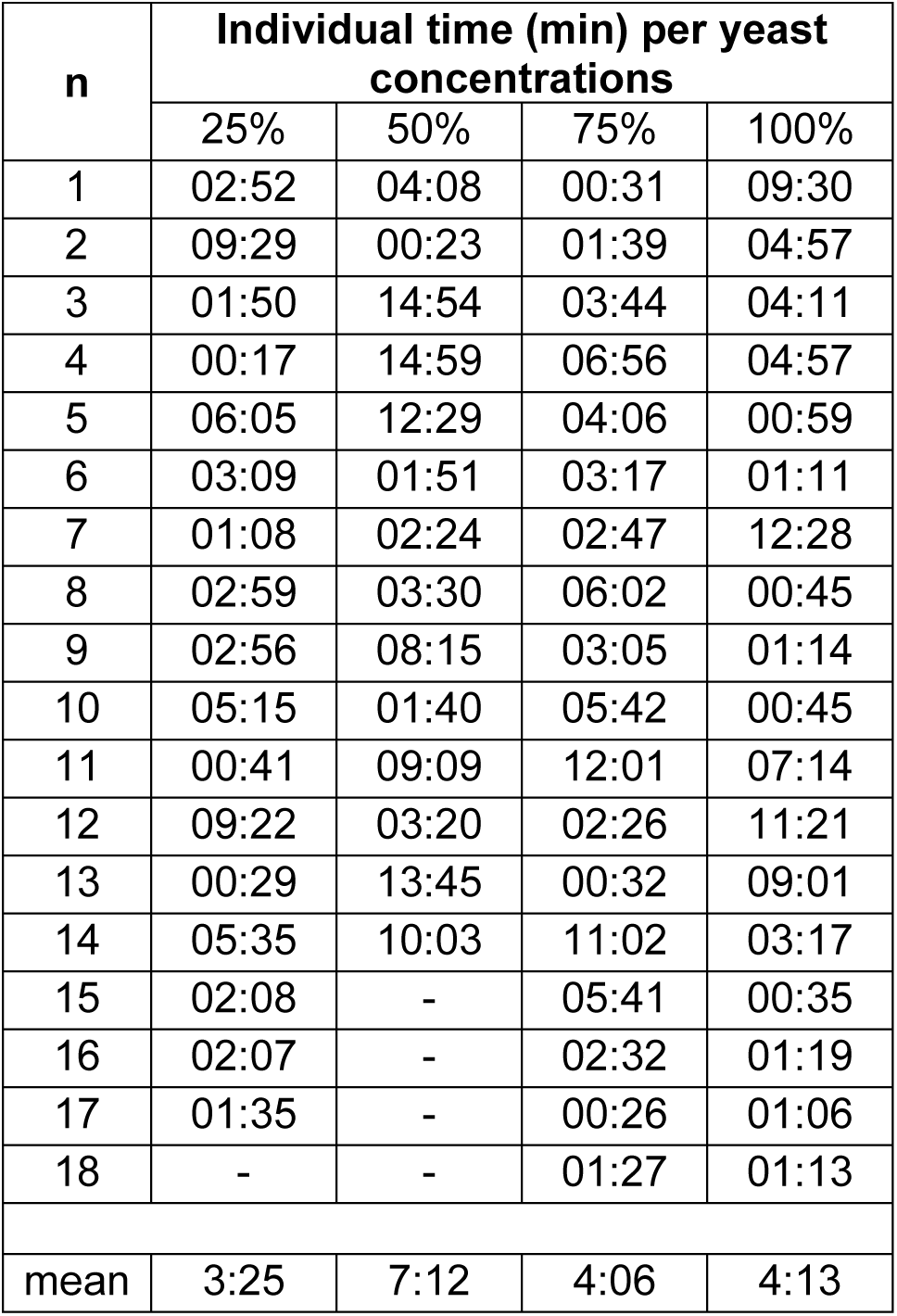
Evaluation of the time larvae spent to find the container. The table shows the time required by the individual larvae to find the container for four different yeast concentrations (25%, 50%, 75% and 100%).

### The influence of the agarose substrate concentration on the local search behavior

Having established a chemical stimulus sufficient to initiate local search behavior, we investigated the impact of additional environmental factors. To assess whether different agarose concentrations affect local search behavior, we presented yeast paste (25%) not only on 0.8% agarose containing (Fig.5A-F, same data as in Fig.4A-F), but also on 1.4% (Fig.5A’-F’) and 2.0% (Fig.5A’’-F’’) agarose containing substrates. All three groups increase their proportion of time and track length within their center-proximate distance categories (Fig.5A-E) and decrease the values of these parameters in the outermost distance categories, resulting in significant increases in their search scores (Fig.5F-F’’). As the agarose concentration increases from 0.8%, to 1.4%, and 2.0%, the time the larvae spend searching in the center decreases (Fig.5D-D’’). Nevertheless, there are significant differences in the local search behavior. During the *baseline phase* the larvae tested on 1.4% and 2.0% containing agarose plates spent more time at the outermost edge (74% and 60% versus 39%). At 0.8% agarose concentration 63.6%, at 1.4% agarose concentration 30.7% and at 2.0% agarose concentration only 8.3% of the larvae were digging into the substrate (data not shown). Therefore, in order to prevent the larvae from digging in, we decided to conduct further experiments with 2.0% agarose substrates.

**Figure 5:**
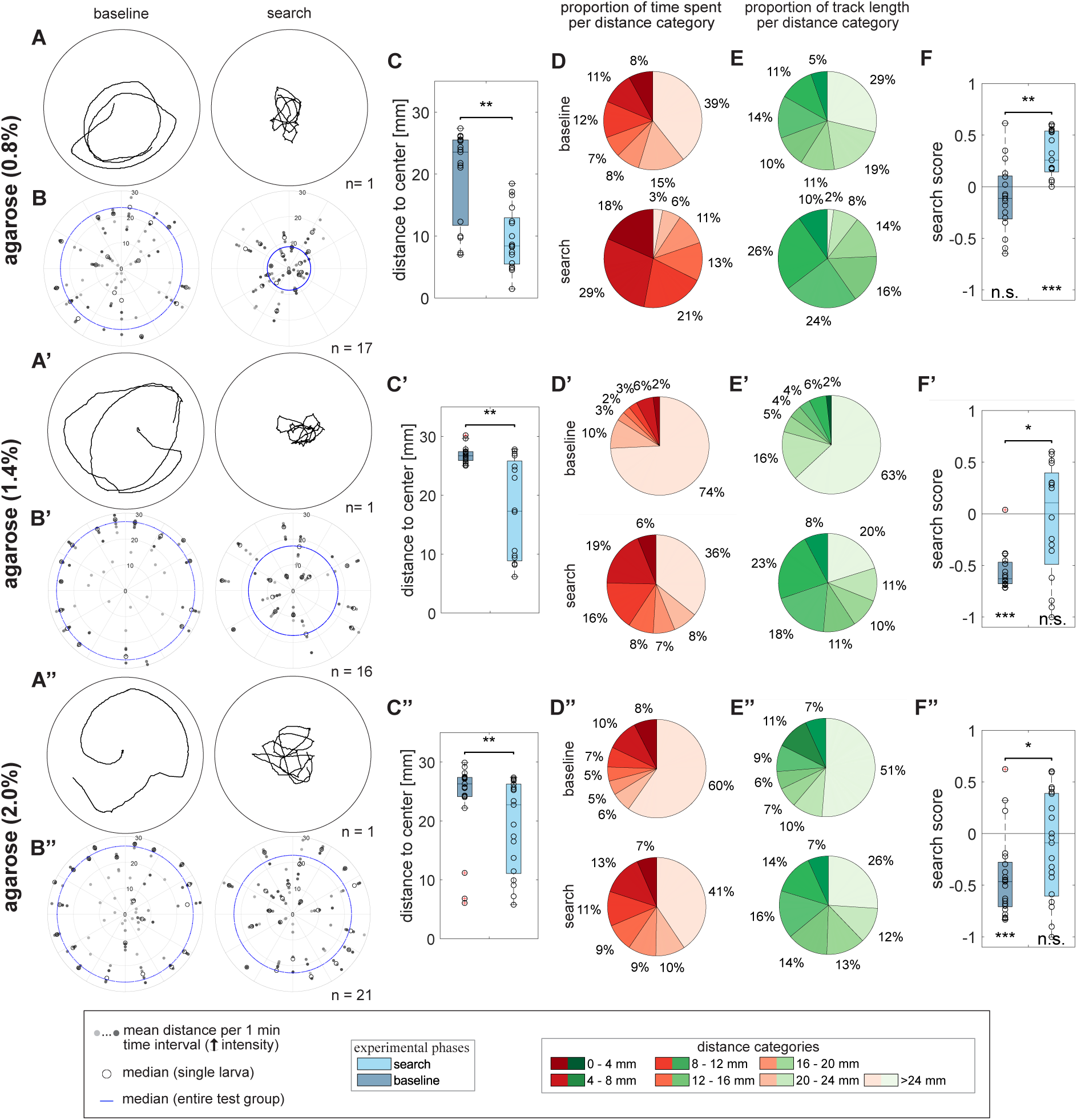
Rising agarose concentrations decrease the local search. Larvae were tested via the local search paradigm to examine the effect of different agarose concentrations (0.8%, 1.4% and 2.0%). (A) Example track. The plot shows the crawled path before and after the presentation of yeast. (B) Distance to center – polar scatter. The figure displays the individual mean distances to center within one-minute time intervals, the individual median distance during the entire phase and the median distance of the entire test group. Individuals are represented by circles on a line extending from the center. (C) Distance to center – boxplot. Compared are the median distances of larvae during the respective phases. All test groups lowered their distance to center significantly after the container interaction (F: p=0.002, F’: p=0.003, F’’: p=0.006). (D) Proportion of time spent per distance category. Pie charts represent the proportion of time spent per distance category counterclockwise from the center (dark red) to the edge (beige). (E) Proportion of track length crawled per distance category. Pie charts represent the proportion of track length crawled per distance category counterclockwise from the center (darkest green) to the edge (lightest green). (F) Search score. Larvae tested on 0.8% agarose plates neither preferred nor avoided the search zone before the stimulus presentation (F: p=0.210) but showed a preference towards it after the interaction (F: p=0.003). The remaining test groups avoided the search zone during the *baseline phase* (F’: p<0.001; F’’: p<0.001) and shifted into a neutral one after the stimulus presentation (F’: p=0.717; F’’: p=0.296). All larvae increased their search score significantly (F: p=0.003, F’: p=0.011, F’’: p=0.042). Larvae were tested after one-hour starvation using yeast (25%). Notice that the data of the 0.8% agarose test group are taken from the yeast (25%) experiment and were therefore not run in parallel to the other agarose concentrations. For the statistical evaluation the one-sample and two-sample Wilcoxon signed-rank test were performed. *p≤0.05, **p≤0.01, ***p<0.001

### Fermented yeast enhances the larval local search

To further enhance the observed search behavior, we used fermented yeast as a stimulus. In a separate experiment, we covered the lid of a container with fermented yeast to investigate the impact of additional gustatory stimulation. Larvae exposed to fermented yeast exhibited a distinct local search behavior, evident in a noticeable reduction of their distance to the center after interacting with the container compared to the *baseline phase* (Fig.6A-C). This effect was even stronger when an additional gustatory stimulus was added to the lid (Fig.6A’-C’). In both cases larvae reduced their time spent and track length crawled within the outermost distance and increased these parameter values within the categories closest to the center (Fig.6D-E,D’-E’). Larvae that had consumed actual food remained within the reward zone for more than half of the time (53%) and covered 43% of their total track length in that area. Conversely, larvae without food intake spent only 19% of their time in the reward zone but traversed 24% of their total track length. Notably, these larvae showed a significant increase in speed after exposure to fermented yeast (Fig.S2, Movie 3; middle and right). The larval search score also revealed the behavioral shift of the two groups (Fig.6F,F’).

**Figure 6:**
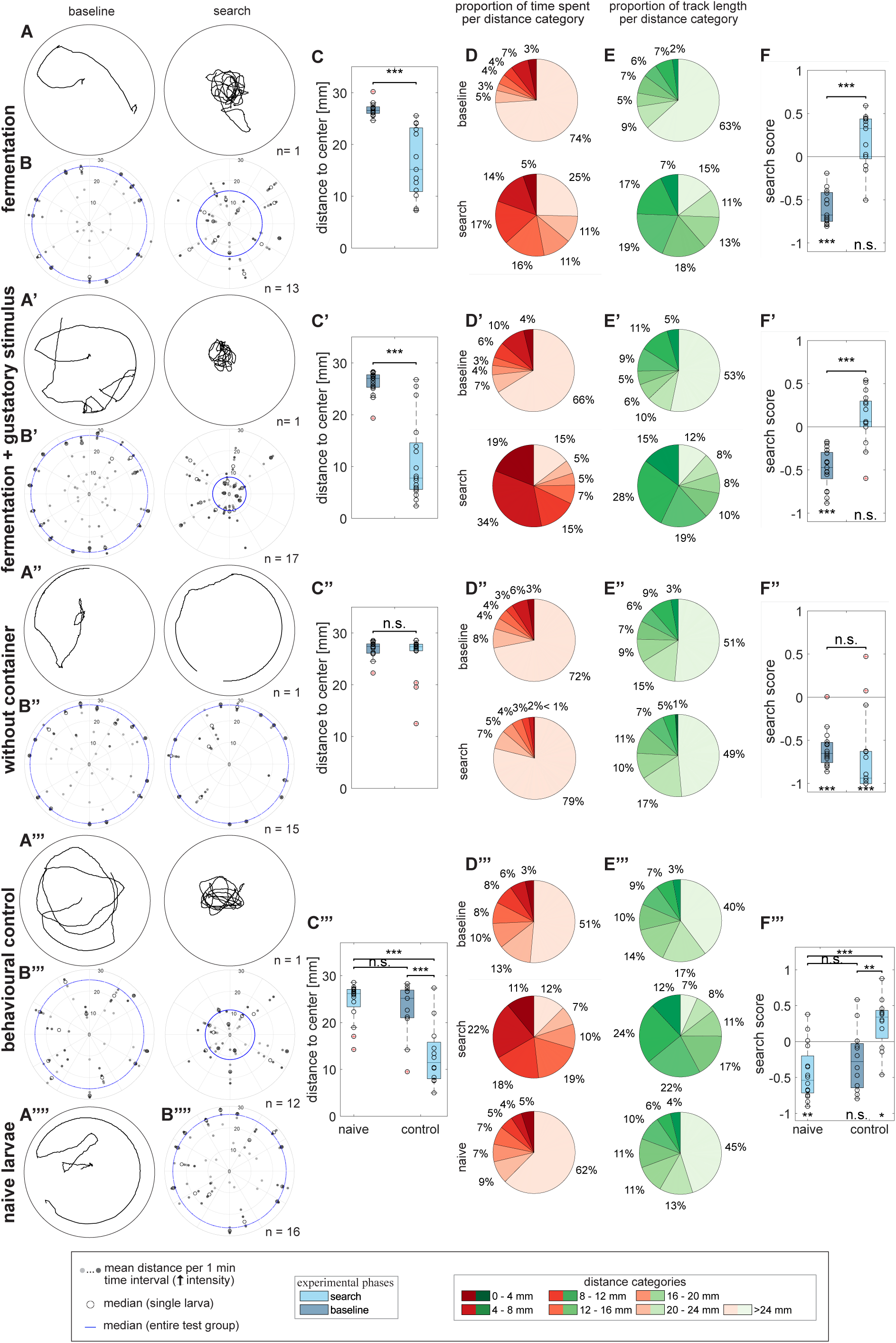
Neither temporal effects nor potential lingering chemical cues affect the local search. Larvae were tested via the local search paradigm to investigate the effect of fermentation, gustatory intake (side walls of the container wetted with fermented yeast), time (without container) and potentially remaining odor cues (naïve larvae). (A) Example track. The plot shows the crawled path before and after the presentation of a chemical stimulus (except without container, here: no stimulus). (B) Distance to center – polar scatter. The figure displays the individual larval mean distances to center within one-minute time intervals, the individual median distance during the entire phase and the median distance of the entire test group. Individual larvae are represented by circles on a line extending from the center. (C) Distance to center – boxplot. Compared are the median distance of larvae during the respective phases. Larvae tested with fermented yeast only or fermented yeast with an additional gustatory stimulus reduced their distance to center after the container interaction (C: p<0.001, C’: p<0.001). Larvae did not reduce their distance to center over time (C’’: p=0.340). The tests with naïve animals showed that behavioral controls reduced their distance to center after the container interaction while naïve animals showed a *baseline*-like behavior (C’’’: p_control_<0.001, p_naïve-base_=0.593, p_naïve-search_<0.001). (D) Proportion of time spent per distance category. Pie charts represent the proportion of time spent per distance category counterclockwise from the center (dark red) to the edge (beige). (E) Proportion of track length crawled per distance category. Pie charts represent the proportion of track length crawled per distance category counterclockwise from the center (darkest green) to the edge (lightest green). (F) Search score. Larvae tested with fermented yeast only or fermented yeast with an additional gustatory stimulus preferred the edge before the stimulus was presented (F: p<0.001; F’: p<0.001) while increasing their search score significantly (F: p<0.001, F’: p<0.001) toward a neutral behavior after the container interaction (F: p=0.080; F’: p=0.119). Without an external stimulus, larvae avoided the center during the *baseline* and *search phase* (F’’: p_base_<0.001, p_search_<0.001) and, therefore, did not change their search score significantly (F’’: p=0.309). Naïve larvae avoided the search zone during the test (F’’’: p_naïve_=0.043) while the behavioral control behaved neutrally during the *baseline phase* and preferred the center after the container interaction (F’’’: p_base_=0.064, p_search_=0.043). The *baseline phase* of behavioral control animals is indistinguishable from the behavior of naïve animals, but both differ from the *search phase* of behavioral control animals (F’’’: p_naïve-base_=0.330, p_naïve-search_<0.001, p_base-search_ =0.005). Larvae were tested after one-hour starvation on 2.0% agarose plates. Naïve larvae were placed after a stimulus propagation time of 7:16 min (mean time the larvae needed to find the container during fermentation experiments). The behavioral controls and naïve larvae were tested by using fermented yeast. For the statistical evaluation the one-sample and two-sample Wilcoxon signed-rank test were performed. For the comparison of naïve larvae with the behavioral control the Mann Whitney U test was used. *p≤0.05, **p≤0.01, ***p<0.001

Finally, we conducted two additional control experiments. The first involved a control without the use of a container (Fig.6A’’-F’’). Throughout the *baseline* and *search phases* the larvae exhibited identical movement patterns, maintained consistent distances to the center, spent the same time in the same areas and moved the same track lengths therein. As a result, their search scores remained unchanged. In the second experiment, naïve larvae were placed at the beginning of the *search phase* on a plate that previously contained a container filled with fermented yeast. The naïve animals showed *baseline*-like behavior compared to a control group running in parallel. They remained at greater distances from the center (Fig.6C’’’) and therefore showed also comparable low search scores (Fig.6F’’’). The proportion of time spent, and the length of the track covered per distance category was also more similar to that of the *baseline phase* of the controls (Fig.6D’’’’/E’’’’, see Fig.S3). In summary, the findings imply that the larvae do not reduce their distance to the center over time or in response to potential lingering chemical stimuli. However, since we cannot quantify the amount of food intake per larva or the presence of yeast remaining, we used the fermented yeast without gustatory input as the chemical stimulus for subsequent experiments.

### The duration of larval starvation affects the local search behavior

In the subsequent experiment, we examined whether the local search behavior is influenced by their feeding state. We compared the local search behavior of well-fed larvae with those that had been starved for three and six hours. Regardless of their starvation status, all tested groups exhibited local search behavior during the *search phase* (see Fig.7A-A’’,B-B’’,C-C’’). However, the reduction in distance to the center diminished with prolonged hunger time yet remained distinct for all test groups between the *baseline* and *search phases*. The distribution of time spent and the distance covered per distance category showed similarities across the individual test groups (see Fig.7D-D’’,E-E’’). Notably, all test groups displayed significant changes in search scores (see Fig.7F-F’’). Given the more consistent and highly significant values observed in experiments with one-hour-starved larvae (Fig.6F), we defined this parameter as standard for the experiment.

**Figure 7:**
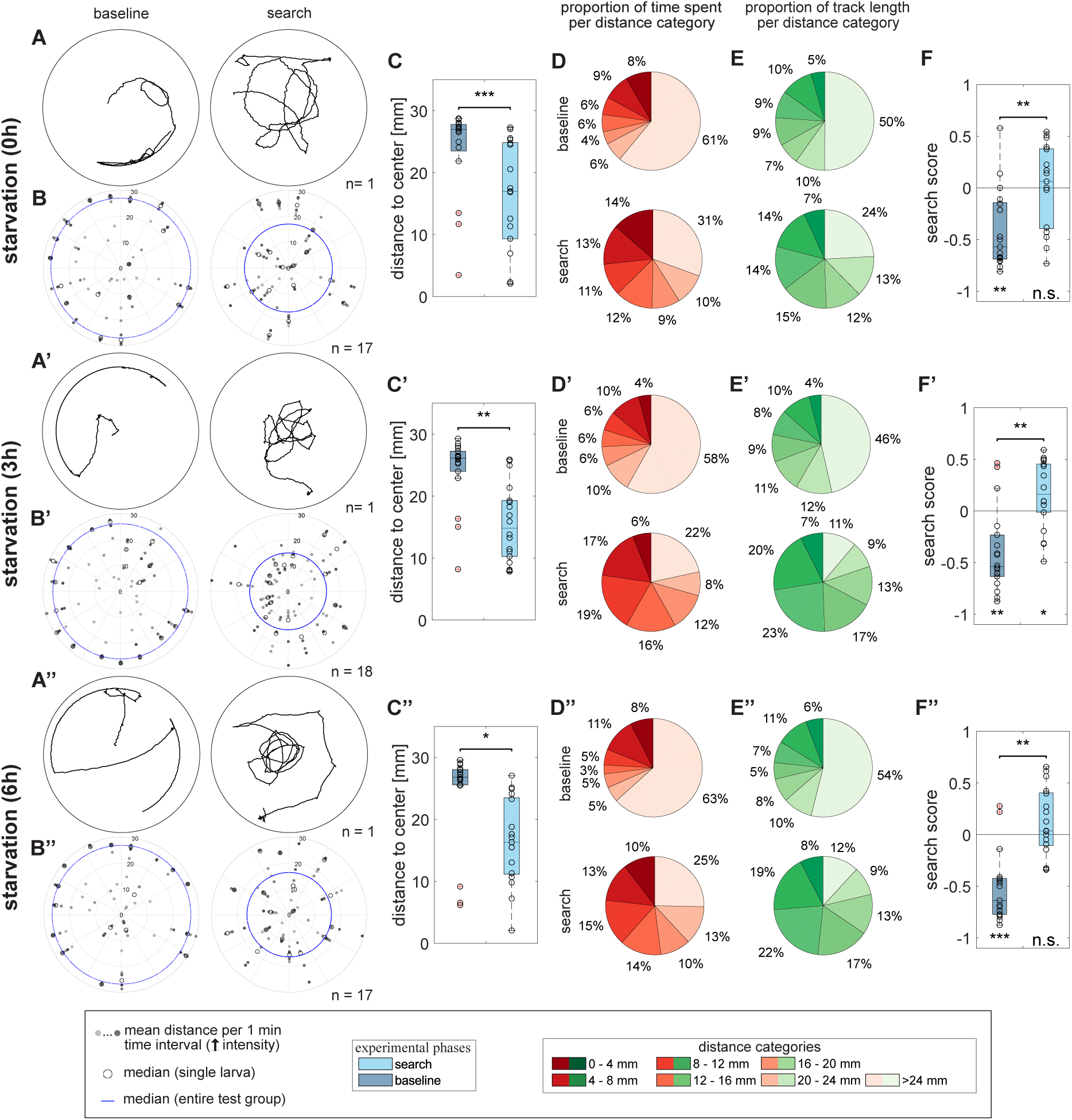
Effect of starvation time on the local search. Larvae were tested via the local search paradigm to investigate the effect of starvation time (0h, 3h, 6h) on the local search. (A) Example track. The plot shows the crawled path before and after the presentation of fermented yeast. (B) Distance to center – polar scatter. The figure displays the individual mean distances to center within one-minute time intervals, the individual median distance during the entire phase, and the median distance of the entire test group. Individuals are represented by circles on a line extending from the center. (C) Distance to center – boxplot. Compared are the median distance of larvae during the respective phases. All test groups reduced their distance to center significantly after the container interaction (C: p<0.001, C’: p=0.006, C’’: p=0.017). (D) Proportion of time spent per distance category. Pie charts represent the proportion of time spent per distance category counterclockwise from the center (dark red) to the edge (beige). (E) Proportion of track length crawled per distance category. Pie charts represent the proportion of track length crawled per distance category counterclockwise from the center (darkest green) to the edge (lightest green). (F) Search score. All test groups preferred the edge before the odor container was presented and only preferred the search zone during the *search phase* after a starvation time of three hours (F: p_base_=0.003, p_search_=0.847; F’: p_base_=0.003, p_search_=0.039; F’’: p_base_<0.001, p_search_=0.196). However, an increasing search score was observed for all the three test groups (F: p=0.001, F’: p=0.003, F’’: p=0.002). The larvae were tested on 2.0% agarose plates using fermented yeast as stimulus. For the statistical evaluation the one-sample and two-sample Wilcoxon signed-rank test were performed. *p≤0.05, **p≤0.01, ***p<0.001

### Larvae perform local search behavior independent of their genetic background

Due to our previous experiments, we defined the following parameters for the larval local search standard experiment: 1 h starvation time, fermented yeast as chemical stimulus, and an agarose concentration of 1.4% for the substrate.

Next, we evaluated the impact of the genetic background on the local search. In all the experiments described thus far, larvae with a specific mutation in the *white* gene (*w^1118^*) were used. This mutation was chosen due to its widespread use as a genetic background in most available fly lines. Therefore, we have replicated the behavior observed in *w^1118^* mutants using wild-type Canton S (*WT-CS*) larvae. Both test groups showed nearly identical behavior. The larvae resided in edge-near distances during the *baseline phase* and showed a highly significant edge preference (Fig.8A,A’,B,B’,C,C’,F,F’). Following the interaction with the container, larvae positioned themselves significantly closer to the center and strongly reduced the time spent within the edge of the arena (66%→16% for *w^1118^*, 62%→13% for *WT-CS*, Fig.8D,D’). The same applies for the track length crawled at the edge (Fig.8E,E’). Calculated search scores confirmed the similar behavior of the two groups. We therefore conclude that the observed behavior in both, *w^1118^* and *WT-CS* larvae, is comparable, enabling future studies on diverse mutant and genetically modified animals.

**Figure 8:**
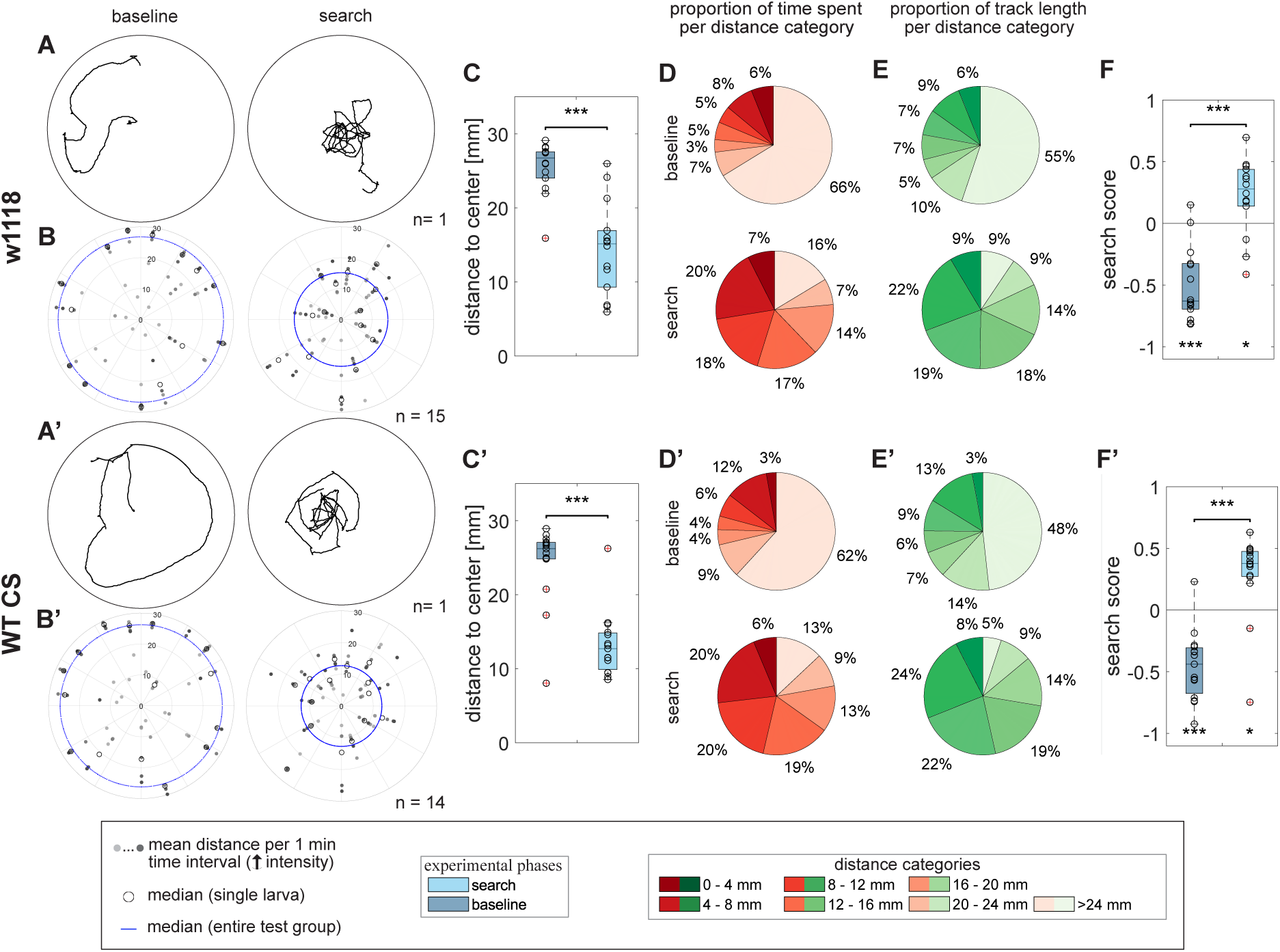
Influence of the genetic background on the local search. To test whether the genetic background of *w^1118^* impacts the search behavior we compared them with *WT-CS*. (A) Example track. The plot shows the crawled path before and after the presentation of fermented yeast. (B) Distance to center – polar scatter. The figure displays the individual mean distances to center within one-minute time intervals, the individual median distance during the entire phase, and the median distance of the entire test group. Individuals are represented by circles on a line extending from the center. (C) Distance to center – boxplot. Compared are the median distance of larvae during the respective phases. In comparison to the *baseline phase*, both test groups remained closer to the center after the container was removed (C: p<0.001, C’: p<0.001). (D) Proportion of time spent per distance category. Pie charts represent the proportion of time spent per distance category counterclockwise from the center (dark red) to the edge (beige). (E) Proportion of track length moved per distance category. Pie charts represent the proportion of track length crawled per distance category counterclockwise from the center (darkest green) to the edge (lightest green). (F) Search score. Both test groups avoided the center during the *baseline phase* (F: p<0.001; F’: p<0.001). After the container was removed, both test groups preferred the search area (F: p=0.025; F’: p=0.010) and, therefore, increased their search score significantly (F: p<0.001, F’: p<0.001). Larvae were tested after a starvation time of one-hour on 1.4% agarose plates using fermented yeast as stimulus. For the statistical evaluation the one-sample and two-sample Wilcoxon signed-rank test were performed. *p≤0.05, **p≤0.01, ***p<0.001.

## Discussion

### *Drosophila* larvae initially show centrophobism/thigmotaxis

We developed the local search paradigm to study larval navigation to a previous food source in an almost featureless habitat. The search behavior is triggered by a chemosensory stimulus, which is subsequently removed. Naïve larvae placed in the center of an arena without an additional stimulus leave the middle within 30 seconds and remain at the edge (Fig.1). At low agarose concentrations, which the larvae prefer, they can also dig in the middle of the plate to protect themselves from predators and dehydration (Kudow et al., 2019). Adult flies show a similar behavior, as they usually avoid central areas in an arena - either as naïve animals or induced after ether anesthesia (Besson and Martin, 2005; Götz and Biesinger, 1985). This behavior is referred to as centrophobism and/or thigmotaxis and is reminiscent of the open field test in rodents (Archer, 1973). Centrophobism/thigmotaxis is also seen in larval and adult zebrafish and other insects like cockroaches, earwigs, ants and larval honeybees (Durier and Rivault, 2003; Kalueff et al., 2013; Laurent Salazar et al., 2018; Perttunen, 1952; Schnorr et al., 2012; Stringer et al., 2017; Vázquez and Farina, 2021). In adult *Drosophila* avoidance of the center has been diminished in adult learning mutants (*dunce*) or after chemical ablation of the MB (Götz and Biesinger, 1985). Here, specifically the MB γ-lobe neurons seems to be necessary (Besson and Martin, 2005). These neurons are already functional in the embryonic and larval stages and form a specific lobe system in the adult animal after they have completely remodeled their morphology during metamorphosis (Armstrong et al., 1998; Technau and Heisenberg, 1982; Truman et al., 2023). It is therefore tempting to speculate that the initially observed larval local search behavior might also require the involvement of the larval MB.

### Yeast is a potent stimulus to trigger local search behavior

Upon the brief presentation of a food stimulus, larval behavior undergoes a notable shift, characterized by the suppression of centrophobism/thigmotaxis and increased movement towards the center where the food source was located (Fig.1). Complex odors emanating from food sources such as apple juice or yeast initiate local search behavior, in contrast to the single odor AM (Fig.3), a key aromatic compound found in fruits. AM is attractive to larvae in olfactory choice assays and effective as a conditioned stimulus in classical odor conditioning assays (Chen and Gerber, 2014; Cobb and Dannet, 1994; Pauls et al., 2010; Scherer et al., 2003; Widmann et al., 2016). Variations in yeast concentrations do not significantly impact the intensity of the induced local search behavior (Fig.4). Yet, larvae display a more centered local search behavior when exposed to fermented yeast odor or when provided with fermented yeast for consumption (Fig.6). This robust response is somehow expected, given that yeast is the principal food source for both larvae and adults of many *Drosophila* species. Yeast not only supplies essential nutrients but also enhances the availability or mitigates the toxicity of certain compounds, thereby influencing larval growth, survival, and body size (Grangeteau et al., 2018). The specific odors associated with *Drosophila* attraction to yeast ferments remain rather unclear. Highly fermenting *Saccharomyces* species, which typically dominate ripe fruit fermentation, produce volatile compounds such as ethanol, volatile acids, aldehydes, ethyl esters, ethyl phenols, and acetate esters, which are generally attractive to larvae (Günther and Goddard, 2019; Schumann et al., 2021). The fly’s olfactory system is equipped with highly conserved ORs to detect these metabolites. Or9a and Or92a detect acetoin, Or42b detects ethyl acetate, Or71a detects ethyl phenols, and Or67a and Or85d detect phenyl ethanol and phenylethyl acetate, respectively (Becher et al., 2010; Dweck et al., 2015; Stokl et al., 2010). Direct chemosensory detection of glycerol, a sugar alcohol produced by yeast, is mediated by the gustatory receptor Gr64a (Wisotsky et al., 2011). However, this gustatory receptor is not expressed in larvae. Therefore, larvae most likely use an alternative taste pathway potentially mediated by ionotropic receptors Ir76b and Ir25a-expressing neurons that respond to multiple stimuli provided by the yeast (Steck et al., 2018).

### Local search behavior in Drosophila

A more detailed parametric analysis revealed that starvation had only a minor influence on search behavior (Fig.7). It increases a little and becomes more stable after approximately one hour of starvation, but then decreases slightly with prolonged starvation. However, also fed larvae clearly show a local search behavior (Fig.7). This contrasts with adult *Drosophila* and blowflies that perform a sugar-elicited local search behavior that is dependent on their starvation status (Bell et al., 1985; Dethier, 1957; Kim and Dickinson, 2017; Murata et al., 2017). Adult *Drosophila* are often starved for about one day in these kinds of experiments. Larvae, due to their high metabolic rate and rapid growth, may have a generally higher hunger level, which quickly results in negative effects when food is further limited, compounded by the high baseline energy demands of the organism. A similar effect is observed in appetitive classical conditioning experiments, which requires adults but not larvae to be food deprived (Gruber et al., 2013; Krashes and Waddell, 2008).

Regarding the behavior itself, differences are evident when comparing adults and larvae. Adults move in short, straight segments interrupted by saccadic turns, often returning to the exact location of a distant food source (Behbahani et al., 2021; Corfas et al., 2019; Kim and Dickinson, 2017; Murata et al., 2017). In contrast, larvae tend to follow a rough circular path around the initial food stimulus and often do not return to the exact starting point (Fig.1,2). This difference may stem from distinct strategies each must employ. Recent experiments show that flies possess head direction cells in the CX that help maintain an absolute sense of orientation (Seelig and Jayaraman, 2015). Such cells have not been identified in larvae, and the reconstruction of the larval connectome does not yet reveal any obvious corresponding wiring pattern (Winding et al., 2023). Consequently, it is likely that the simpler cellular structure of the larval brain does not support this function, resulting in less precise and less position-accurate local search behavior.

### Larval search strategies

What search strategy could the larva use? We would like to mention four different options in ascending complexity: simple changes of the motor pattern, sensory taxis, place memory, and idiothetic path integration. A straightforward explanation for local foraging behavior could be that the brief food stimulus shifts the larvae’s movement pattern from an exploratory mode to a more local one. A similar fundamental shift in foraging behavior is reported for the foraging gene; larvae with the rover allele travel longer distances than those with the sitter allele (Osborne et al., 1997; Sokolowski, 1980). These different foraging patterns are achieved by altering the speed and frequency of pauses and turns (Gomez-Marin et al., 2011; Sokolowski, 1980). In the simplest scenario, the yeast stimulus slows down the larvae’s movement and increases the number of pauses for several minutes, keeping them closer to the center of the arena (Wosniack et al., 2022). However, the larvae in our assay behave in the opposite way. During the search, they increase their basic speed in response to fermented yeast (Fig.2K). Therefore, other strategies must be present.

Larvae might also receive sensory cues during its local search that cause it to turn back toward the center as it moves away from it, as the gradient of sensory stimuli decreases. Consequently, the larva engages in a directed movement, or taxis. But what could this sensory stimulus be? Larvae possess a simple visual system that is incapable of perceiving infrared light, which we utilized in our experiments, allowing us to rule out visual stimuli (Humberg et al., 2018; Keene and Sprecher, 2012; Sprecher et al., 2011). Likewise, no specific acoustic or tactile stimuli were present in the experimental setup that the larva could use. We also minimized the potential influence of taste by enclosing the yeast and apple juice in a container with a perforated lid so that no food residuals could get onto the agarose. Therefore, we propose that the only remaining means for the larva to orient itself is through olfactory cues. This could be due to lingering residues of yeast and apple juice odors, or pheromones that the larvae use to mark the location of the food source (Mast et al., 2014). However, when naïve larvae are placed on test plates where previously fermented yeast was presented, they do not exhibit local search behavior (Fig.6). In addition, the odor stimulus from the container is very weak. For apple juice, only a good third of the larvae even found the container in the arena. Likewise, the direct contact time is limited to only one minute, so it is unlikely that this time is sufficient to set a pheromone signal. Therefore, this explanation does not seem completely convincing either.

It is also possible that the larvae develop a spatial memory for the position of the stimulus. Social insects like bees and ants use memorized visual cues to maintain direction (alignment image-matching) and navigate to familiar locations (positional image-matching) (Collett et al., 2013). Similarly, adult *Drosophila* can recognize and remember a visual environment in a heat maze test to find a cool spot in a heated arena (Ofstad et al., 2011). They can also recall the direction of a visual stimulus even when it is no longer present and remember the spatial location of a cold spot in a warm environment using mechanosensory cues (Neuser et al., 2008; Ostrowski et al., 2015; Wustmann et al., 1996). Thus, larvae might employ a comparable form of memory. Mechanosensory information might be at play. Therefore, one could now investigate which mechanosensory neurons or brain areas are crucial for the local search behavior.

Finally, larvae might use idiothetic rather than allothetic cues to locate a previously visited feeding site, similar to strategies observed in adult flies and other insects (Corfas et al., 2019; Kim and Dickinson, 2017; Muller and Wehner, 1988; Wittlinger et al., 2006). While larvae have sensory neurons capable of detecting proprioceptive feedback and internal copies of motor commands, they lack a CX, which is essential for tracking distances and orientations during movement and for using this information to determine their current position (Agrawal and Tuthill, 2022; Greaney et al., 2023; Richter et al., 2024; Vaadia et al., 2019; Winding et al., 2023). Yet, larvae display highly stereotyped locomotor behavior, characterized by the synchronous contraction of muscles. During forward crawling, peristaltic waves of muscle contractions move from the posterior to the anterior abdominal segments, while the opposite occurs during backward crawling. During turns, the unilateral contraction of one side of the body causes the front part to swing from side to side (head casting). The direction of a turn is determined by this casting behavior, which ends with the front part of the body at an angle to the rest of the body (Berni et al., 2012; Dixit et al., 2008; Fox et al., 2006; Gjorgjieva et al., 2013; Heckscher et al., 2012; Lahiri et al., 2011). As the larva moves forward, the hind part gradually reorients to align with the front. This rhythmic movement, consisting of runs and turns, can be generated independently of sensory feedback and descending input from the brain, suggesting the presence of a central pattern generator network in the thoracic and abdominal segments (Berni et al., 2012; Hughes and Thomas, 2007; Suster and Bate, 2002). Given its simplicity, this raises the possibility of a basic crawl counter mechanism, similar in concept to a step counter observed in ants.

With this study providing the first parametric description of the larva’s local search behavior, it should now be possible to decipher the underlying neuronal network in the next step. Since this foraging behavior is not unique to *Drosophila* larvae, it will also be intriguing to determine whether a rudimentary larval CX exists - or if the larva has developed a different coding strategy.

## Supporting information

all supplemental information

## List of symbols and abbreviations colcnt

AM: amyl acetate
Crv: center revisits
CX: central complex
EB: ellipsoid body
E-PG: ellipsoid body-protocerebral bridge-gall neurons
FB: fan-shaped body
GRN: gustatory receptor neuron
MB: mushroom body
NO: noduli
OR: olfactory receptor
ORN: olfactory receptor neuron
PB: protocerebral bridge
WT-CS: wild type Canton-S

## Supplemental Information

Supplemental Information includes Supplemental Experimental Procedures and additional Figures and can be found with this article online.

## Acknowledgments

This work was supported by the Deutsche Forschungsgemeinschaft (Grant No. 441181781, 426722269, 432195391) and by EU funds from the ESF Plus Program (Grant No. 100649752) all to AST. We thank Bert Klagges, Dennis Pauls, Mareike Selcho, and Wolf Huetteroth for discussions and comments. Additionally, we thank Juliane Saumweber for fly care and maintenance.

## Author contributions

Conceptualization, J.K., T.T, and A.S.T; Methodology, J.K., T.T., and A.S.T; Investigation, J.K; Writing, J.K., T.T., and A.S.T; Supervision, A.S.T.

## References

Agrawal, S. and Tuthill, J. C. (2022). The two-body problem: Proprioception and motor control across the metamorphic divide. Curr Opin Neurobiol 74, 102546.

Archer, J. (1973). Tests for emotionality in rats and mice: a review. Anim Behav 21, 205–35.

Armstrong, J. D., de Belle, J. S., Wang, Z. and Kaiser, K. (1998). Metamorphosis of the mushroom bodies; large-scale rearrangements of the neural substrates for associative learning and memory in Drosophila. Learn Mem 5, 102–14.

Becher, P. G., Bengtsson, M., Hansson, B. S. and Witzgall, P. (2010). Flying the fly: long-range flight behavior of Drosophila melanogaster to attractive odors. J Chem Ecol 36, 599–607.

Behbahani, A. H., Palmer, E. H., Corfas, R. A. and Dickinson, M. H. (2021). Drosophila re-zero their path integrator at the center of a fictive food patch. Curr Biol 31, 4534–4546 e5.

Bell, W. J. (1990). Searching Behavior Patterns in Insects. Annual Review of Entomology 35, 447–467.

Bell, W. J., Cathy, T., Roggero, R. J., Kipp, L. R. and Tobin, T. R. (1985). Sucrose-stimulated searching behaviour of Drosophila melanogaster in a uniform habitat: modulation by period of deprivation. Animal Behaviour 33, 436–448.

Berni, J., Pulver, S. R., Griffith, L. C. and Bate, M. (2012). Autonomous circuitry for substrate exploration in freely moving Drosophila larvae. Curr Biol 22, 1861–70.

Besson, M. and Martin, J.-R. (2005). Centrophobism/thigmotaxis, a new role for the mushroom bodies in Drosophila. Journal of Neurobiology 62, 386–396.

Bures, J., Fenton, A. A., Kaminsky, Y., Wesierska, M. and Zahalka, A. (1998). Rodent navigation after dissociation of the allocentric and idiothetic representations of space. Neuropharmacology 37, 689–99.

Chen, Y. C. and Gerber, B. (2014). Generalization and discrimination tasks yield concordant measures of perceived distance between odours and their binary mixtures in larval Drosophila. J Exp Biol 217, 2071–7.

Cobb, M. and Dannet, F. (1994). Multiple genetic control of acetate-induced olfactory responses in Drosophila melanogaster larvae. Heredity (Edinb) 73 (Pt 4), 444–55.

Collett, M., Chittka, L. and Collett, Thomas S. (2013). Spatial Memory in Insect Navigation. Current Biology 23, R789–R800.

Corfas, R. A., Sharma, T. and Dickinson, M. H. (2019). Diverse Food-Sensing Neurons Trigger Idiothetic Local Search in Drosophila. Curr Biol 29, 1660–1668 e4.

Degen, J., Kirbach, A., Reiter, L., Lehmann, K., Norton, P., Storms, M., Koblofsky, M., Winter, S., Georgieva, P. B., Nguyen, H. et al. (2016). Honeybees Learn Landscape Features during Exploratory Orientation Flights. Curr Biol 26, 2800–2804.

Dethier, V. G. (1957). Communication by Insects: Physiology of Dancing. Science 125, 331–6.

Dixit, R., Vijayraghavan, K. and Bate, M. (2008). Hox genes and the regulation of movement in Drosophila. Dev Neurobiol 68, 309–16.

Durier, V. and Rivault, C. (2003). Exploitation of home range and spatial distribution of resources in German cockroaches (Dictyoptera: Blattellidae). J Econ Entomol 96, 1832–7.

Dweck, H. K., Ebrahim, S. A., Farhan, A., Hansson, B. S. and Stensmyr, M. C. (2015). Olfactory proxy detection of dietary antioxidants in Drosophila. Curr Biol 25, 455–66.

Eichler, K., Li, F., Litwin-Kumar, A., Park, Y., Andrade, I., Schneider-Mizell, C. M., Saumweber, T., Huser, A., Eschbach, C., Gerber, B. et al. (2017). The complete connectome of a learning and memory centre in an insect brain. Nature 548, 175–182.

Eschbach, C., Fushiki, A., Winding, M., Afonso, B., Andrade, I. V., Cocanougher, B. T., Eichler, K., Gepner, R., Si, G., Valdes-Aleman, J. et al. (2021). Circuits for integrating learned and innate valences in the insect brain. Elife 10.

Farnworth, M. S., Eckermann, K. N. and Bucher, G. (2020). Sequence heterochrony led to a gain of functionality in an immature stage of the central complex: A fly-beetle insight. PLoS Biol 18, e3000881.

Fishilevich, E., Domingos, A. I., Asahina, K., Naef, F., Vosshall, L. B. and Louis, M. (2005). Chemotaxis behavior mediated by single larval olfactory neurons in Drosophila. Curr Biol 15, 2086–96.

Fox, L. E., Soll, D. R. and Wu, C. F. (2006). Coordination and modulation of locomotion pattern generators in Drosophila larvae: effects of altered biogenic amine levels by the tyramine beta hydroxlyase mutation. J Neurosci 26, 1486–98.

Gendre, N., Luer, K., Friche, S., Grillenzoni, N., Ramaekers, A., Technau, G. M. and Stocker, R. F. (2004). Integration of complex larval chemosensory organs into the adult nervous system of Drosophila. Development 131, 83–92.

Gershow, M., Berck, M., Mathew, D., Luo, L., Kane, E. A., Carlson, J. R. and Samuel, A. D. (2012). Controlling airborne cues to study small animal navigation. Nat Methods 9, 290–6.

Giraldo, Y. M., Leitch, K. J., Ros, I. G., Warren, T. L., Weir, P. T. and Dickinson, M. H. (2018). Sun Navigation Requires Compass Neurons in Drosophila. Curr Biol 28, 2845–2852 e4.

Gjorgjieva, J., Berni, J., Evers, J. F. and Eglen, S. J. (2013). Neural circuits for peristaltic wave propagation in crawling Drosophila larvae: analysis and modeling. Front Comput Neurosci 7, 24.

Gomez-Marin, A., Stephens, G. J. and Louis, M. (2011). Active sampling and decision making in Drosophila chemotaxis. Nat Commun 2, 441.

Götz, K. G. and Biesinger, R. (1985). Centrophobism inDrosophila melanogaster. Journal of Comparative Physiology A 156, 319–327.

Grangeteau, C., Yahou, F., Everaerts, C., Dupont, S., Farine, J. P., Beney, L. and Ferveur, J. F. (2018). Yeast quality in juvenile diet affects Drosophila melanogaster adult life traits. Sci Rep 8, 13070.

Greaney, M. R., Wreden, C. C. and Heckscher, E. S. (2023). Distinctive features of the central synaptic organization of Drosophila larval proprioceptors. Frontiers in Neural Circuits 17.

Green, J., Adachi, A., Shah, K. K., Hirokawa, J. D., Magani, P. S. and Maimon, G. (2017). A neural circuit architecture for angular integration in Drosophila. Nature 546, 101–106.

Green, J., Vijayan, V., Mussells Pires, P., Adachi, A. and Maimon, G. (2019). A neural heading estimate is compared with an internal goal to guide oriented navigation. Nat Neurosci 22, 1460–1468.

Gruber, F., Knapek, S., Fujita, M., Matsuo, K., Bracker, L., Shinzato, N., Siwanowicz, I., Tanimura, T. and Tanimoto, H. (2013). Suppression of conditioned odor approach by feeding is independent of taste and nutritional value in Drosophila. Curr Biol 23, 507–14.

Günther, C. S. and Goddard, M. R. (2019). Do yeasts and Drosophila interact just by chance? Fungal Ecology 38, 37–43.

Hanesch, U., Fischbach, K. F. and Heisenberg, M. (1989). Neuronal architecture of the central complex in Drosophila melanogaster. Cell and Tissue Research 257, 343–366.

Heckscher, E. S., Lockery, S. R. and Doe, C. Q. (2012). Characterization of Drosophila larval crawling at the level of organism, segment, and somatic body wall musculature. J Neurosci 32, 12460–71.

Hughes, C. L. and Thomas, J. B. (2007). A sensory feedback circuit coordinates muscle activity in Drosophila. Mol Cell Neurosci 35, 383–96.

Humberg, T. H., Bruegger, P., Afonso, B., Zlatic, M., Truman, J. W., Gershow, M., Samuel, A. and Sprecher, S. G. (2018). Dedicated photoreceptor pathways in Drosophila larvae mediate navigation by processing either spatial or temporal cues. Nat Commun 9, 1260.

Kalueff, A. V., Gebhardt, M., Stewart, A. M., Cachat, J. M., Brimmer, M., Chawla, J. S., Craddock, C., Kyzar, E. J., Roth, A., Landsman, S. et al. (2013). Towards a comprehensive catalog of zebrafish behavior 1.0 and beyond. Zebrafish 10, 70–86.

Keene, A. C. and Sprecher, S. G. (2012). Seeing the light: photobehavior in fruit fly larvae. Trends Neurosci 35, 104–10.

Kim, I. S. and Dickinson, M. H. (2017). Idiothetic Path Integration in the Fruit Fly Drosophila melanogaster. Curr Biol 27, 2227–2238 e3.

Krashes, M. J. and Waddell, S. (2008). Rapid consolidation to a radish and protein synthesis-dependent long-term memory after single-session appetitive olfactory conditioning in Drosophila. J Neurosci 28, 3103–13.

Kudow, N., Kamikouchi, A. and Tanimura, T. (2019). Softness sensing and learning in Drosophila larvae. J Exp Biol 222.

Lahiri, S., Shen, K., Klein, M., Tang, A., Kane, E., Gershow, M., Garrity, P. and Samuel, A. D. (2011). Two alternating motor programs drive navigation in Drosophila larva. PLoS One 6, e23180.

Larsson, M. C., Domingos, A. I., Jones, W. D., Chiappe, M. E., Amrein, H. and Vosshall, L. B. (2004). Or83b encodes a broadly expressed odorant receptor essential for Drosophila olfaction. Neuron 43, 703–14.

Laurent Salazar, M.-O., Planas-Sitjà, I., Sempo, G. and Deneubourg, J.-L. (2018). Individual Thigmotactic Preference Affects the Fleeing Behavior of the American Cockroach (Blattodea: Blattidae). Journal of Insect Science 18.

Mast, J. D., De Moraes, C. M., Alborn, H. T., Lavis, L. D. and Stern, D. L. (2014). Evolved differences in larval social behavior mediated by novel pheromones. Elife 3, e04205.

Menzel, R., Geiger, K., Chittka, L., Joerges, J., Kunze, J., Uuml and Ller, U. (1996). The knowledge base of bee navigation. J Exp Biol 199, 141–6.

Mittelstaedt, M. L. and Mittelstaedt, H. (2001). Idiothetic navigation in humans: estimation of path length. Exp Brain Res 139, 318–32.

Muller, M. and Wehner, R. (1988). Path integration in desert ants, Cataglyphis fortis. Proc Natl Acad Sci U S A 85, 5287–90.

Murata, S., Brockmann, A. and Tanimura, T. (2017). Pharyngeal stimulation with sugar triggers local searching behavior in Drosophila. J Exp Biol 220, 3231–3237.

Neuser, K., Triphan, T., Mronz, M., Poeck, B. and Strauss, R. (2008). Analysis of a spatial orientation memory in Drosophila. Nature 453, 1244–7.

Ofstad, T. A., Zuker, C. S. and Reiser, M. B. (2011). Visual place learning in Drosophila melanogaster. Nature 474, 204–7.

Osborne, K. A., Robichon, A., Burgess, E., Butland, S., Shaw, R. A., Coulthard, A., Pereira, H. S., Greenspan, R. J. and Sokolowski, M. B. (1997). Natural behavior polymorphism due to a cGMP-dependent protein kinase of Drosophila. Science 277, 834–6.

Ostrowski, D., Kahsai, L., Kramer, E. F., Knutson, P. and Zars, T. (2015). Place memory retention in Drosophila. Neurobiol Learn Mem 123, 217–24.

Pauls, D., Selcho, M., Gendre, N., Stocker, R. F. and Thum, A. S. (2010). Drosophila larvae establish appetitive olfactory memories via mushroom body neurons of embryonic origin. J Neurosci 30, 10655–66.

Perttunen, V. (1952). Seasonal Change in the Humidity Reaction of the Common Earwig, Forficula auricularia. Nature 170, 209–210.

Python, F. and Stocker, R. F. (2002). Adult-like complexity of the larval antennal lobe of D. melanogaster despite markedly low numbers of odorant receptor neurons. J Comp Neurol 445, 374–87.

Richter, V., Rist, A., Kislinger, G., Laumann, M., Schoofs, A., Miroschnikow, A., Pankratz, M., Cardona, A. and Thum, A. S. (2024). Morphology and ultrastructure of external sense organs of Drosophila larvae: eLife Sciences Publications, Ltd.

Riebli, N., Viktorin, G. and Reichert, H. (2013). Early-born neurons in type II neuroblast lineages establish a larval primordium and integrate into adult circuitry during central complex development in Drosophila. Neural Dev 8, 6.

Risse, B., Thomas, S., Otto, N., Lopmeier, T., Valkov, D., Jiang, X. and Klambt, C. (2013). FIM, a novel FTIR-based imaging method for high throughput locomotion analysis. PLoS One 8, e53963.

Rist, A. and Thum, A. S. (2017). A map of sensilla and neurons in the taste system of drosophila larvae. J Comp Neurol 525, 3865–3889.

Saumweber, T., Rohwedder, A., Schleyer, M., Eichler, K., Chen, Y. C., Aso, Y., Cardona, A., Eschbach, C., Kobler, O., Voigt, A. et al. (2018). Functional architecture of reward learning in mushroom body extrinsic neurons of larval Drosophila. Nat Commun 9, 1104.

Scherer, S., Stocker, R. F. and Gerber, B. (2003). Olfactory learning in individually assayed Drosophila larvae. Learn Mem 10, 217–25.

Schnorr, S. J., Steenbergen, P. J., Richardson, M. K. and Champagne, D. L. (2012). Measuring thigmotaxis in larval zebrafish. Behav Brain Res 228, 367–74.

Schumann, I., Berger, M., Nowag, N., Schafer, Y., Saumweber, J., Scholz, H. and Thum, A. S. (2021). Ethanol-guided behavior in Drosophila larvae. Sci Rep 11, 12307.

Schumann, I. and Triphan, T. (2020). The PEDtracker: An Automatic Staging Approach for Drosophila melanogaster Larvae. Front Behav Neurosci 14, 612313.

Seelig, J. D. and Jayaraman, V. (2015). Neural dynamics for landmark orientation and angular path integration. Nature 521, 186–91.

Singh, R. N. and Singh, K. (1984). Fine structure of the sensory organs of Drosophila melanogaster Meigen larva (Diptera : Drosophilidae). International Journal of Insect Morphology and Embryology 13, 255–273.

Sokolowski, M. B. (1980). Foraging strategies of Drosophila melanogaster: a chromosomal analysis. Behav Genet 10, 291–302.

Sprecher, S. G., Cardona, A. and Hartenstein, V. (2011). The Drosophila larval visual system: high-resolution analysis of a simple visual neuropil. Dev Biol 358, 33–43.

Steck, K., Walker, S. J., Itskov, P. M., Baltazar, C., Moreira, J. M. and Ribeiro, C. (2018). Internal amino acid state modulates yeast taste neurons to support protein homeostasis in Drosophila. Elife 7.

Stokl, J., Strutz, A., Dafni, A., Svatos, A., Doubsky, J., Knaden, M., Sachse, S., Hansson, B. S. and Stensmyr, M. C. (2010). A deceptive pollination system targeting drosophilids through olfactory mimicry of yeast. Curr Biol 20, 1846–52.

Stringer, L. D., Corn, J. E., Sik Roh, H., Jiménez-Pérez, A., Manning, L.-A. M., Harper, A. R. and Suckling, D. M. (2017). Thigmotaxis Mediates Trail Odour Disruption. Scientific Reports 7, 1670.

Suster, M. L. and Bate, M. (2002). Embryonic assembly of a central pattern generator without sensory input. Nature 416, 174–8.

Technau, G. and Heisenberg, M. (1982). Neural reorganization during metamorphosis of the corpora pedunculata in Drosophila melanogaster. Nature 295, 405–7.

Titova, A. V., Kau, B. E., Tibor, S., Mach, J., Vo-Doan, T. T., Wittlinger, M. and Straw, A. D. (2023). Displacement experiments provide evidence for path integration in Drosophila. J Exp Biol 226.

Truman, J. W., Price, J., Miyares, R. L. and Lee, T. (2023). Metamorphosis of memory circuits in Drosophila reveals a strategy for evolving a larval brain. Elife 12.

Vaadia, R. D., Li, W., Voleti, V., Singhania, A., Hillman, E. M. C. and Grueber, W. B. (2019). Characterization of Proprioceptive System Dynamics in Behaving Drosophila Larvae Using High-Speed Volumetric Microscopy. Current Biology 29, 935–944.e4.

Vafidis, P., Owald, D., D’Albis, T. and Kempter, R. (2022). Learning accurate path integration in ring attractor models of the head direction system. Elife 11.

Vázquez, D. E. and Farina, W. M. (2021). Locomotion and searching behaviour in the honey bee larva depend on nursing interaction. Apidologie 52, 1368–1386.

Weber, D., Richter, V., Rohwedder, A., Grossjohann, A. and Thum, A. S. (2023). Learning and Memory in Drosophila Larvae. Cold Spring Harb Protoc 2023, pdb top107863.

Widmann, A., Artinger, M., Biesinger, L., Boepple, K., Peters, C., Schlechter, J., Selcho, M. and Thum, A. S. (2016). Genetic Dissection of Aversive Associative Olfactory Learning and Memory in Drosophila Larvae. PLoS Genet 12, e1006378.

Winding, M., Pedigo, B. D., Barnes, C. L., Patsolic, H. G., Park, Y., Kazimiers, T., Fushiki, A., Andrade, I. V., Khandelwal, A., Valdes-Aleman, J. et al. (2023). The connectome of an insect brain. Science 379, eadd9330.

Wisotsky, Z., Medina, A., Freeman, E. and Dahanukar, A. (2011). Evolutionary differences in food preference rely on Gr64e, a receptor for glycerol. Nat Neurosci 14, 1534–41.

Wittlinger, M., Wehner, R. and Wolf, H. (2006). The ant odometer: stepping on stilts and stumps. Science 312, 1965–7.

Wolff, T. and Rubin, G. M. (2018). Neuroarchitecture of the Drosophila central complex: A catalog of nodulus and asymmetrical body neurons and a revision of the protocerebral bridge catalog. J Comp Neurol 526, 2585–2611.

Wosniack, M. E., Festa, D., Hu, N., Gjorgjieva, J. and Berni, J. (2022). Adaptation of Drosophila larva foraging in response to changes in food resources. Elife 11.

Wustmann, G., Rein, K., Wolf, R. and Heisenberg, M. (1996). A new paradigm for operant conditioning of Drosophila melanogaster. J Comp Physiol A 179, 429–36.

Zeil, J. (2023). Visual navigation: properties, acquisition and use of views. J Comp Physiol A Neuroethol Sens Neural Behav Physiol 209, 499–514.

Zjacic, N. and Scholz, M. (2022). The role of food odor in invertebrate foraging. Genes Brain Behav 21, e12793.

