## Supplementary material for "Finding a path: Local search behavior of *Drosophila* larvae": all supplemental information

**Supplemental Data**

**Figure S1: Effect of different apple juice concentrations on the local search behavior.** Larvae were tested via the larval local search paradigm to evaluate the influence of different apple juice concentrations, 25%, 50%, 75% and 100%, on the larval behavior.

(A) Example track. The plot shows the traveled path before and after the presentation of the apple juice stimulus.

(B) Distance to center – polar scatter. The figure displays mean distances to center of individual larvae within one-minute time intervals the individual median distance during the entire phase and the median distance of the entire test group. Individual larvae are represented by circles on a line extending from the center.

(C) Distance to center – boxplot. Compared are the median distances of larvae during the *baseline* and *search phase*. Larvae exposed to low (25%), and high (100%) concentrations of apple juice remained closer to the center after the container interaction than before (C: $p=0.019$; C’’’: $p<0.001$). In contrast the distance to center after a presentation of 50% and 75% apple juice is unaffected (C’: $p=0.113$; C’’: $p=0.064$).

(D) Proportion of time spent per distance category. Pie charts represent the proportion of time spent per distance category counterclockwise from the center (dark red) to the edge (beige).

(E) Proportion of track length crawled per distance category. Pie charts represent the proportion of track length crawled per distance category counterclockwise from the center (darkest green) to the edge (lightest green).

(F) Search score. The search score displays whether larvae prefer the search (positive values) or edge (negative values) zone. All test groups preferred the edge during the *baseline phase* and showed a neutral behavior after the container interaction (F: $p_{base}=0.003$, $p_{search}=0.852$; F’: $p_{base}<0.001$, $p_{search}=0.093$; F’’: $p_{base}=0.002$, $p_{search}=0.241$; F’’’:

$p_{base}$=0.001, $p_{search}$=0.776). Only larvae exposed to 25% and 100% apple juice increased their search score significantly (F: p=0.014; F': p=0.163; F'': p=0.489; F''': p=0.016).

The larvae were tested after one-hour starvation time on 0.8% agarose plates. To compare other apple juice concentrations with apple juice (100%), we present parts of the data of Fig.1 and 2 here again. For the statistical evaluation the one-sample and two-sample Wilcoxon signed-rank test were performed. *p≤0.05, **p≤0.01, ***p<0.001

**Figure S2: Impact of potentially lingering chemical cues on the local search behavior and the impact of fermentation products on the larval velocity.** Shown are the results of the larval local search paradigm for naïve larvae placed into the search arena after 15 min of odor evaporation compared to a parallel running behavioral control group (A-F) and the comparison of velocity for larvae tested before and after the presentation of fermented yeast with or without a gustatory stimulus.

(A) Example track. The plot shows the crawled path before and after the presentation of a chemical stimulus.

(B) Distance to center – polar scatter. The figure displays the individual larval mean distances to center within one-minute time intervals the individual median distance during the entire phase and the median distance of the entire test group Circles that are represented in a line display a single larva.

(C) Distance to center – boxplot. Compared are the median distances of larvae during the *baseline* and *search phase*. Naïve larvae performed a *baseline*-like behavior characterized by a high distance to center (p=0.763). Both, naïve larvae, and the behavioral controls *baseline phase*, differ significantly from the search phase ($p_{naïve\text{-}search}$<0.001, $p_{base\text{-}search}$=<0.001).

(D) Proportion of time spent per distance category. Pie charts represent the proportion of time spent per distance category counterclockwise from the center (dark red) to the edge (beige).

(E) Proportion of track length crawled per distance category. Pie charts represent the proportion of track length crawled per distance category counterclockwise from the center (darkest green) to the edge (lightest greens).

(F) Search score. The search score displays whether larvae prefer the search (positive values) or edge (negative values) zone. Naïve larvae preferred the edge zone during their test phase while the behavioral control larvae were neutral during the *baseline phase* and preferred the search zone during their *search phase* ($p_{naïve}$=0.015, $p_{base}$=0.064, $p_{search}$=0.043). The behavior of the naïve animals is indistinguishable from the controls *baseline* but differs from their *search phase* ($p_{naïve-base}$=0.562, $p_{naïve-search}$=0.003, $p_{base-search}$=0.005).

(G) Velocity – boxplot. The boxplot compares the larval velocity during the respective phases. Larvae exposed to the fermented yeast without gustatory intake show a higher velocity after the presentation of the stimulus than before (p<0.001). In contrast, the velocity does not rise significantly if the larvae had an additional gustatory intake (p=0.758).

(H) Velocity - progression over time. The graphs display the mean velocity averaged over ten frames of the test group (blue line) as well as the individual velocity (grey lines) over time.

The larvae were tested after one-hour starvation time on 2.0% agarose plates. For the statistical evaluation the one-sample and two-sample Wilcoxon signed-rank test were performed. *p≤0.05, **p≤0.01, ***p<0.001

**Figure S3: Larval walking trajectories before and after the stimulus presentation.** The visualization displays the larval behavior under different conditions. Following applied to the different test groups: Apple juice, yeast, empty container, and amyl acetate experiments were performed on 0.8% agarose plates, after a starvation time of one hour with the respective odor. Agarose experiments were conducted on crawling grounds with the respective agarose concentration after a starvation time of one hour using yeast (25%) as chemical stimulus. The fermentation, fermentation with additional gustatory stimulus and without container experiments were performed on 2.0% agarose plates after a starvation time of one hour using the respective chemical stimulus. Naïve Larvae were tested on 2.0% agarose plates after a starvation time of one hour using fermented yeast as chemical stimulus. For the starvation experiments larvae were tested on 2.0% agarose plates using fermented yeast as stimulus. The comparison of $w^{1118}$ and *WT-CS* larvae were conducted on 1.4% agarose plates after a starvation time of one hour using fermented yeast as chemical stimulus.

**Table S1: Statistical evaluation of the larval local search paradigm.** The table visualizes the p-values and medians from the two parameters distance to center and search score of the performed experiments. Significant p-values are highlighted in bold.

**Table S2: Statistical evaluation of the apple juice (100%) experiment.** Visualized are the p-values and medians of the different parameters evaluated in Fig.2. Significant p-values are highlighted in bold.

**Table S3: Statistical evaluation of the velocity data.** Visualized are the p-values and medians of the velocity parameter for the fermentation experiment displayed in S3. Significant p-values are highlighted in bold.

**Movie 1: Larval behavior before and after the presentation of 100% apple juice.** The video shows the behavior of a single larva during the entire local search paradigm.

The video was recorded with two frames per second and playback is sped up by a factor of ten.

**Movie 2: Larval behavior before and after the presentation of amyl acetate.** The video shows the behavior of four larvae during the entire local search paradigm. The video was recorded with two frames per second and playback is sped up by a factor of ten.

**Movie 3: Larval behavior before and after the presentation of a yeast paste (25%) and fermented yeast with and without gustatory intake.** The video shows the behavior of a single larva during the entire local search paradigm exposed to yeast (left video), fermented yeast without (middle video) or with feeding possibility (right video). The videos were recorded with two frames per second and playback is sped up by a factor of ten.

| test group / comparisons | distance to center | | search score | | | | displayed in figure |
|---|---|---|---|---|---|---|---|
| | p-values two-sample Wilcoxon signed rank or Mann Whitney U test (median: baseline \| search [mm]) | | p-values two-sample Wilcoxon signed rank or Mann Whitney U test (median: baseline \| search) | | p-values one-sample Wilcoxon signed rank test (baseline \| search) | | |
| **apple juice (100%)** | **0.019** | (25.39 \| 19.39) | **0.014** | (-0.392 \| 0.152) | **0.003** \| 0.852 | | 2/3/S1 |
| **empty** | **0.022** | (21.25 \| 15.49) | **0.005** | (-0.237 \| 0.347) | 0.064 \| 0.073 | | 3 |
| **amyl acetate (1:250)** | 0.934 | (26.34 \| 26.27) | 0.804 | (-0.460 \| -0.533) | **< 0.001 \| 0.005** | | 3 |
| **yeast paste** | **0.002** | (25.59 \| 11.27) | **0.002** | (-0.338 \| 0.008) | **0.001 \| 0.008** | | 3 |
| **yeast (25%)** | **0.002** | (23.56 \| 8.39) | **0.003** | (-0.113 \| 0.258) | 0.210 \| **< 0.001** | | 4/5 |
| **yeast (50%)** | **< 0.001** | (20.70 \| 9.70) | **0.005** | (0.021 \| 0.408) | **0.021 \| 0.003** | | 4 |
| **yeast (75%)** | **0.007** | (23.31 \| 11.70) | **0.018** | (-0.249 \| 0.316) | **0.025 \| 0.049** | | 4 |
| **yeast (100%)** | **< 0.001** | (24.43 \| 8.74) | **0.001** | (-0.257 \| 0.205) | **0.039 \| 0.003** | | 4 |
| **agarose (1.4%)** | **0.003** | (26.72 \| 17.30) | **0.011** | (-0.630 \| 0.107) | **< 0.001** \| 0.717 | | 5 |
| **agarose (2.0%)** | **0.006** | (26.30 \| 22.72) | **0.042** | (-0.465 \| -0.090) | **< 0.001** \| 0.296 | | 5 |
| **fermentation** | **< 0.001** | (26.56 \| 15.17) | **< 0.001** | (-0.677 \| 0.327) | **< 0.001** \| 0.080 | | 6 |
| **fermentation + gustatory** | **< 0.001** | (27.02 \| 7.76) | **< 0.001** | (-0.472 \| 0.060) | **< 0.001** \| 0.119 | | 6 |
| **without container** | 0.340 | (27.42 \| 16.99) | 0.309 | (-0.652 \| -0.938) | **< 0.001 \| < 0.001** | | 6 |
| **starvation (0h)** | **< 0.001** | (27.01 \| 16.99) | **0.001** | (-0.572 \| 0.062) | **0.003** \| 0.847 | | 7 |
| **starvation (3h)** | **0.006** | (26.11 \| 14.85) | **0.003** | (-0.538 \| 0.162) | **0.003 \| 0.039** | | 7 |
| **starvation (6h)** | **0.017** | (26.79 \| 16.31) | **0.002** | (-0.638 \| 0.037) | **< 0.001** \| 0.196 | | 7 |
| **$w^{1118}$** | **< 0.001** | (26.22 \| 12.71) | **< 0.001** | (-0.629 \| 0.280) | **< 0.001 \| 0.025** | | 8 |
| **WTCS** | **< 0.001** | (26.74 \| 15.13) | **< 0.001** | (-0.438\| 0.377) | **< 0.001 \| 0.010** | | 8 |
| **control (baseline) \| naive** | 0.593 | (25.19 \| 26.21) | 0.330 | (-0.279\| -0.538) | (0.064 \| **0.003**) | | 6 |
| **control (search) \| naive** | **< 0.001** | (11.47 \| 26.21) | **< 0.001** | (0.341 \| -0.538) | (**0.043 \| 0.003**) | | 6 |
| **control (baseline) \| control (search)** | **< 0.001** | (25.19 \| 11.47) | **0.005** | (-0.279 \| 0.341) | (0.064 \| **0.043**) | | 6 |
| **apple juice (75%)** | 0.113 | (26.34 \| 22.34) | 0.163 | (-0.487 \| -0.382) | **< 0.001** \| 0.093 | | S1 |
| **apple juice (50%)** | 0.064 | (26.08 \| 20.10) | 0.489 | (-0.435 \| -0.035) | **0.002** \| 0.241 | | S1 |
| **apple juice (25%)** | **< 0.001** | (26.50 \| 17.18) | **0.016** | (-0.467 \| 0.000) | **0.001** \| 0.776 | | S1 |

| | | | | | | |
|---|---|---|---|---|---|---|
| **control (baseline) \| naive** | 0.763 | (25.19 \| 24.96) | 0.562 | (-0.279 \| -0.343) | (0.064 \| **0.015**) | S2 |
| **control (search) \| naive** | **< 0.001** | (11.48 \| 24.96) | **0.003** | (0.341 \| -0.343) | (**0.043** \| **0.015**) | S2 |
| **control (baseline) \| control (search)** | **< 0.001** | (25.19 \| 11.48) | **0.005** | (-0.279 \| 0.341) | (0.064 \| **0.043**) | S2 |

Table S1

| | **p-values two-sample Wilcoxon signed rank test (median baseline \| search)** |
|---|---|
| **Fig.2D (distance [mm] per time interval)** | |
| **1** | 0.135 (12.18 \| 12.05) |
| **2** | **0.004** (24.47 \| 15.08) |
| **3** | **0.003** (25.07 \| 20.91) |
| **4** | 0.126 (24.79 \| 21.36) |
| **5** | **0.037** (25.12 \| 19.87) |
| **Fig.2E (time spent in reward zone [min])** | |
| | 0.391 (0.58 \| 0.41) |
| **Fig.2F (time spent [s] per distance category)** | |
| **0-4 mm** | 0.446 (6.25 \| 4.25) |
| **4-8 mm** | 0.173 (25.50 \| 24.50) |
| **8-12 mm** | **0.038** (16.00 \| 48.75) |
| **12-16 mm** | **0.018** (12.50 \| 31.25) |
| **16-20 mm** | 0.117 (14.00 \| 21.25) |
| **20-24 mm** | 0.809 (31.50 \| 31.75) |
| **> 24 mm** | **0.014** (190.00 \| 81.50) |
| **Fig.2H (track length [cm])** | |
| | **0.017** (24.13 \| 27.91) |
| **Fig.2I (track length [cm] per distance category)** | |
| **0-4 mm** | 0.396 (0.43 \| 0.51) |
| **4-8 mm** | **0.040** (1.83 \| 2.85) |
| **8-12 mm** | **0.023** (1.67 \| 4.77) |
| **12-16 mm** | **0.015** (1.46 \| 3.43) |
| **16-20 mm** | 0.145 (1.51 \| 2.34) |
| **20-24 mm** | 0.841 (3.35 \| 2.94) |
| **> 24 mm** | **0.010** (12.08 \| 4.27) |
| **Fig.2K (velocity [mm $s^{-1}$])** | |
| | **0.001** (0.64 \| 0.92) |
| **Fig.2M (centre revisits [crv])** | |
| | 0.092 (2.0 \| 2.5) |

Table S2

| test group | p-values two-sample Wilcoxon signed rank test (median: baseline \| search [mm $s^{-1}$]) |
|---|---|
| **fermentation** | **< 0.001** (0.71 \| 1.19) |
| **fermentation + gustatory** | 0.758 (0.66 \| 0.76) |

Table S3

### Figure S1

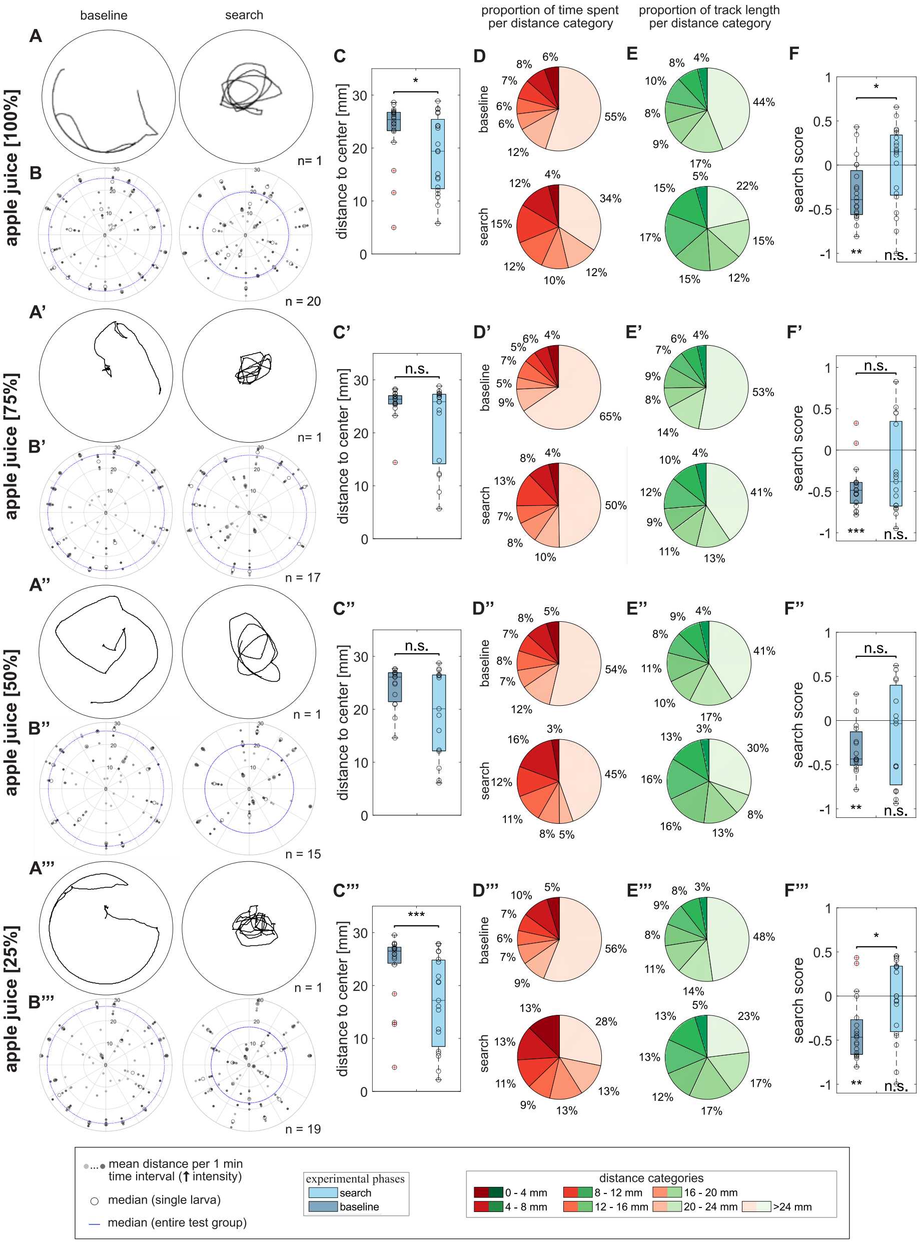

baseline
search
proportion of time spent per distance category
proportion of track length per distance category
apple juice [100%]
apple juice [75%]
apple juice [50%]
apple juice [25%]
distance to center [mm]
search score
n = 20
n = 17
n = 15
n = 19
mean distance per 1 min time interval (↑ intensity)
median (single larva)
median (entire test group)
experimental phases
search
baseline
distance categories
0 - 4 mm
4 - 8 mm
8 - 12 mm
12 - 16 mm
16 - 20 mm
20 - 24 mm
>24 mm

Figure S2

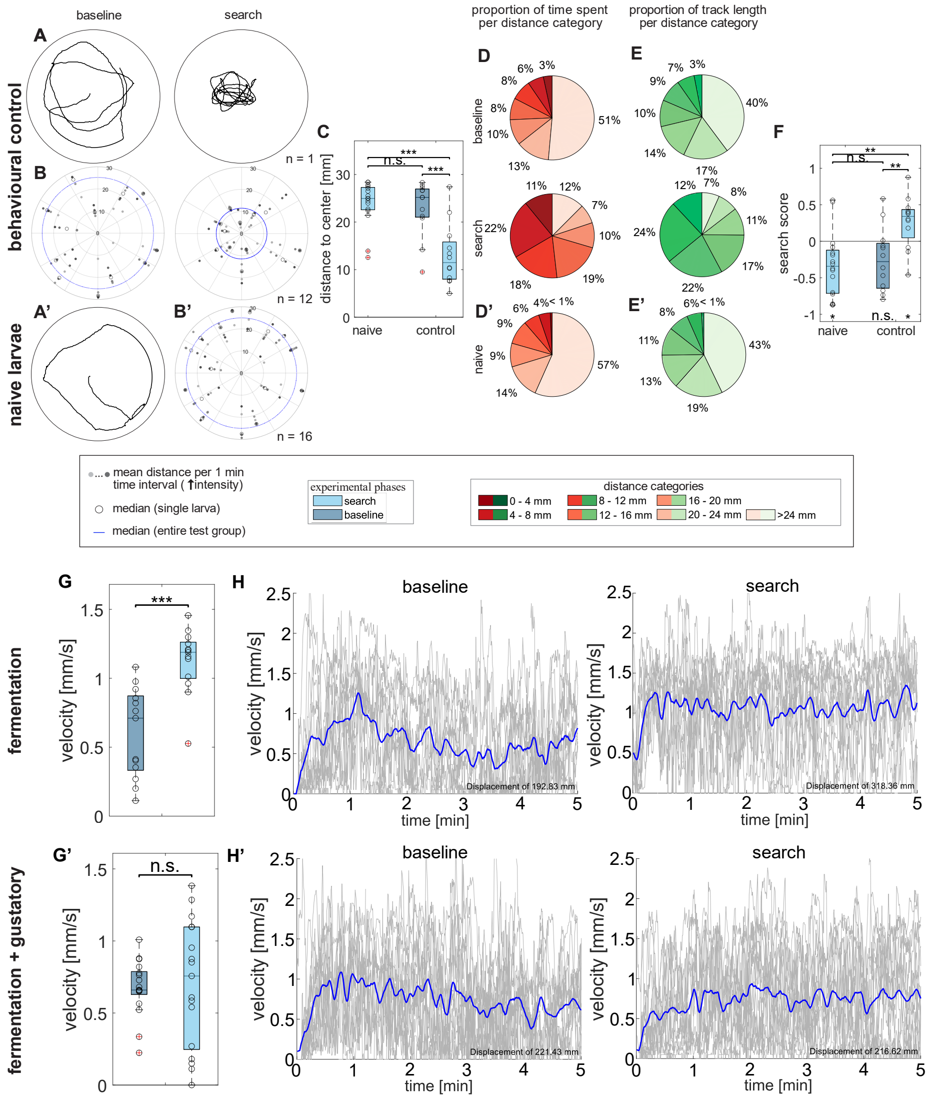
baseline
search
proportion of time spent per distance category
proportion of track length per distance category
behavioural control
naive larvae
n = 1
n = 12
n = 16
distance to center [mm]
naive
control
n.s.
***
search score
mean distance per 1 min time interval (↑intensity)
median (single larva)
median (entire test group)
experimental phases
search
baseline
distance categories
0 - 4 mm
4 - 8 mm
8 - 12 mm
12 - 16 mm
16 - 20 mm
20 - 24 mm
>24 mm
fermentation
fermentation + gustatory
velocity [mm/s]
time [min]
Displacement of 192.83 mm
Displacement of 318.36 mm
Displacement of 221.43 mm
Displacement of 216.62 mm

Figure S3

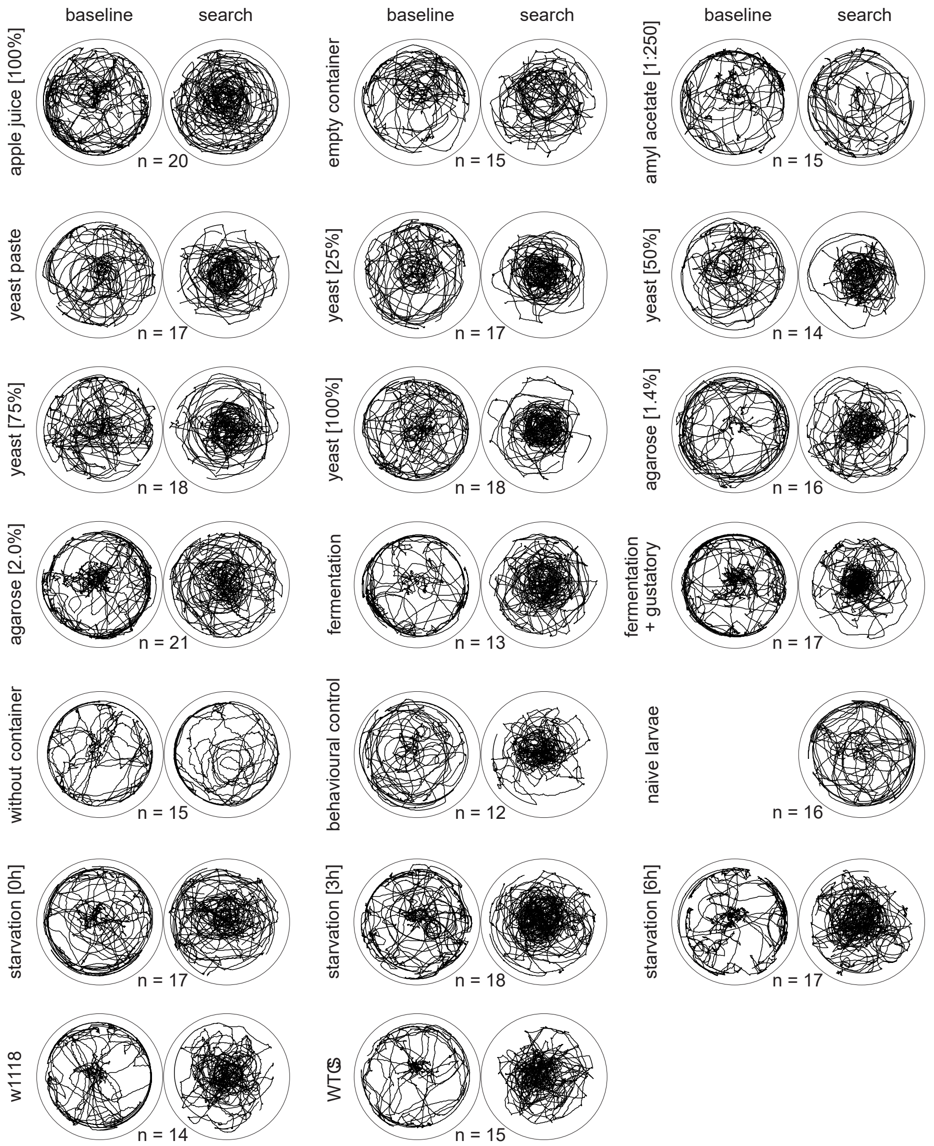
baseline
search
apple juice [100%]
n = 20
empty container
n = 15
amyl acetate [1:250]
n = 15
yeast paste
n = 17
yeast [25%]
n = 17
yeast [50%]
n = 14
yeast [75%]
n = 18
yeast [100%]
n = 18
agarose [1.4%]
n = 16
agarose [2.0%]
n = 21
fermentation
n = 13
fermentation + gustatory
n = 17
without container
n = 15
behavioural control
n = 12
naive larvae
n = 16
starvation [0h]
n = 17
starvation [3h]
n = 18
starvation [6h]
n = 17
w1118
n = 14
WTCS
n = 15
